# Comparative demography elucidates the longevity of parasitic and symbiotic relationships

**DOI:** 10.1101/271965

**Authors:** Luke B.B. Hecht, Peter C. Thompson, Benjamin M. Rosenthal

**Affiliations:** Oak Ridge Institute for Science Education, Oak Ridge, TN; Agricultural Research Service, US Department of Agriculture. 10300 Baltimore Avenue, Beltsville, MD 20705

## Abstract

Parasitic and symbiotic relationships govern vast nutrient and energy flows^1,2,^ but controversy surrounds their longevity. Enduring relationships may engender parallel phylogenies among hosts and parasites^3,4,^ but so may more ephemeral relationships when parasites disproportionately colonize related hosts^5^. When considering these relationships’ temporal durability, it would be useful to understand whether parasite and host populations have grown and contracted in concert. Here, we devised methods to compare demographic histories, derived from genomic data^6^. We used these methods to compare the historical growth of the agent of severe human malaria, *Plasmodium falciparum*, to human and primate histories^7,8^ and to that of their mosquito vector *Anopheles gambiae*^9^, thereby discerning long-term parallels and anthropogenic population explosions^10,11^. The growth history of *Trichinella spiralis*, a zoonotic parasite disseminated by swine domestication^12,13^, proved regionally-specific, paralleling distinctive growth histories for wild boar in Asia and Europe^14^. Parallel histories were inferred for an anemone and its algal symbiont (*Aiptasia pallida^15^ and Symbiodinium minutum*^16^). Concerted growth in potatoes and the agent of potato blight (*Solanum tuberosum^17^ and Phytophthora infestans*^18^) did not commence until the age of potato domestication, helping date the acquisition of this historically consequential fungal plant pathogen. Therefore, comparative historical demography provides a powerful new means by which to interrogate the history of myriad ecological relationships, enriching our understanding of their origins and durability.

---

In order to examine the demographic history of individual species, we determined the extent and distribution of heterozygous positions called from high-quality genome assemblies. Each demographic history was then reconstructed using the pairwise sequentially Markovian coalescent (PSMC) model, a procedure that uses the distribution of heterozygous sites in a diploid genome to estimate the size of the diploid individual’s ancestral population over time^6^. To synchronize such plots, we devised a quantitative curve-fitting procedure to estimate the relative generation time of organismal pairs. We then assessed whether and when ancestral populations grew or contracted in concert. To explore the power of these methods, we tested whether natural species pairs (e.g. malaria and humans) matched each other significantly better than unnatural pairs (e.g. malaria and wild boars (Supplemental Figure S2, S3)); we further established the temporal specificity of this approach, verifying that synchronized demographic plots fit each other significantly better than chronologically disordered comparisons (Supplemental Figure S1). In the case of *P. falciparum*, which is only briefly diploid and which has been sequenced in its haploid state, we used bioinformatic procedures to synthesize “pseudodiploid” chromosomes from isolate pairs (Supplemental Methods).

We found that *P. falciparum* and *A. gambiae* populations grew and shrank in concert with human demography over a ~500K year interval, and not with other great apes (Figure 1A and Supplemental Information). Populations grew steadily, tripling between 500-200 KYA, then declined over the following 170 KY to about 20% their maximum, and stabilized for ~20 KY. Populations then began to grow 3-11 KYA. Notably, these parasites and vectors paralleled the explosive growth unique (among primates) to human beings since the Neolithic, complementing a recent comprehensive survey of anopheline populations^11^. Generation times derived from our curve-fitting procedure closely approximated prior empirical estimates. Assuming 25-year generations for primates, we estimated for *P. falciparum* and *A. gambiae* 2.08 and 17.5 generations/year, respectively (previously reported as 2 and 10-24 generations/year, respectively^10,19,20^). Our findings support prior conclusions that forest clearing and agricultural settlements engendered growth in anopheline populations and intensified their role as vectors of malaria^21^.

**Figure 1:**
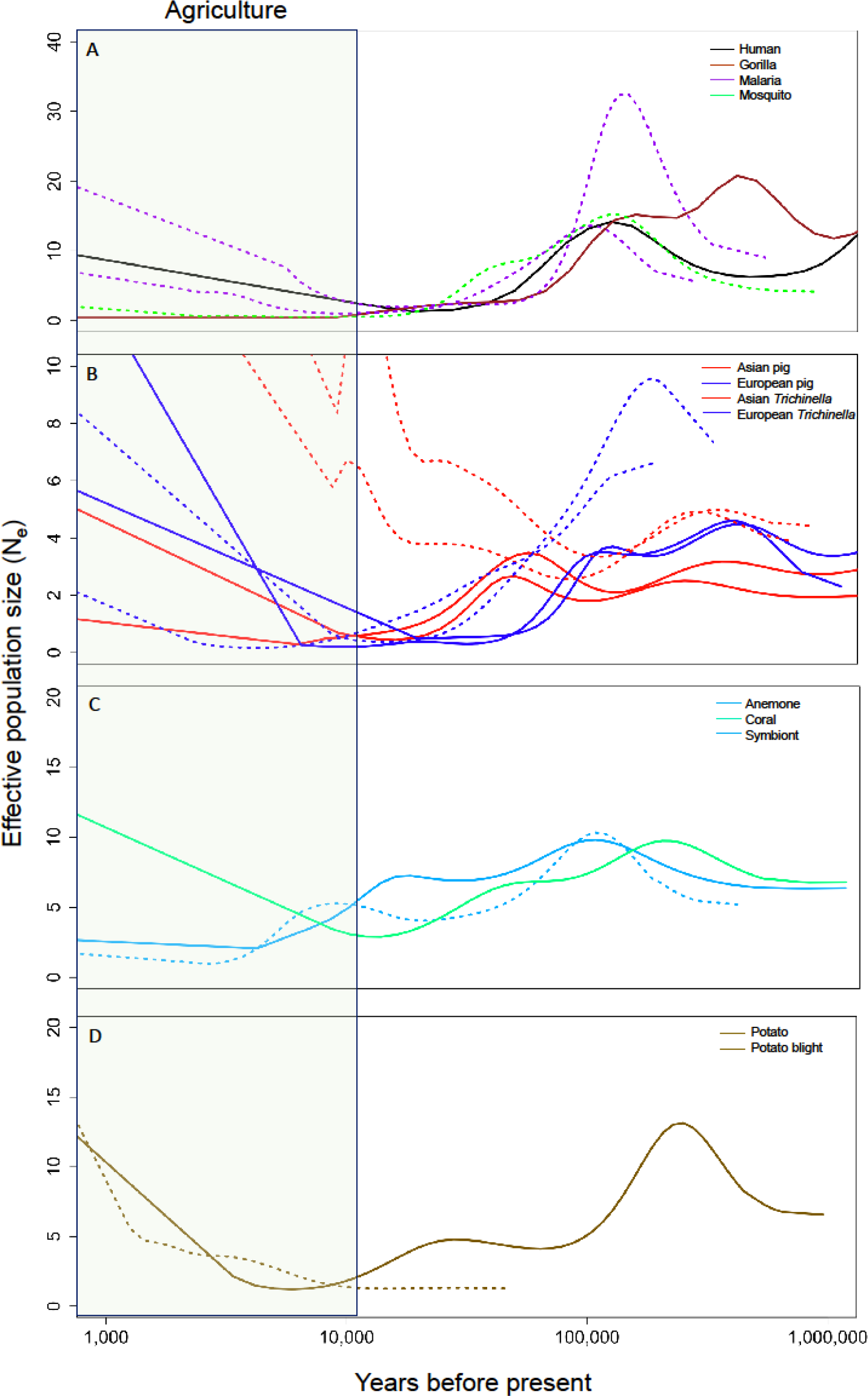
PSMC plots comparing the demographic histories of hosts and their putative parasites/symbionts. The time period associated with human agriculture (most recent 12,000 years) is shaded. Solid lines represent hosts whereas dotted lines represent their parasites or symbionts. A) *A. gambiae* mosquito and *P. falciparum* parasite populations appear to have grown and contracted in sync with human, but not gorilla, populations over the last 300,000 years. Demography estimated from each of two pseudo-diploid malaria genomes illustrated (see Supplemental methods). B) Chinese *T. spiralis* mirror the demographic history of sympatric members of their host species, contrasting with wild boars from Europe. C) The algal symbiont *S. minutum* has tracked the demography of its known host, the anemone *Aiptasia pallida*, rather than the coral *Acropora digitifera*, at least since the LGM (~20 ka). D) *P. infestans*, the agent of potato blight, only shows concerted growth or decline with potatoes since domestication.

Distinct demographic histories have been reported for wild boar in Europe and Asia^14^; notably, Asian *T. spiralis* specifically matches that for sympatric wild boar, growing between 800 KYA and 400 KYA, declining, and then growing 100 KYA-40 KYA; populations of boar in Europe, by contrast, peaked ~100 KYA and declined until ~10 KYA. In Europe and North America, *T. spiralis* are so lacking in heterozygosity (presumably owing to a severe population bottleneck) that PSMC cannot estimate demographic history before 200 KYA (Figure 1B). Nonetheless, contemporaneous comparisons of western *T. spiralis* and European wild boar match significantly better than do asynchronous comparisons (Supplemental Figure S1). Highly reproducible demographies from 100 KYA-10 KYA substantiate explosive growth of *T. spiralis*, in each region, during the last 20 KY (Figure 1 and Supplemental Information). Transmission evidently intensified after the advent of agriculture, likely among abundant (but inbred) domesticated swine and rats.

In spite of climate-driven bleaching events that might have rendered ephemeral the symbiotic relationship between an anemone and its photosynthetic symbiont (*Aiptasia pallida* and *S. minutum*), strikingly similar and temporally-specific demographic histories were recovered (Figure 1 and Supplemental Information). During and since the last glacial maximum (LGM), ancestors in the Red Sea experienced common constraints. Prior to the LGM, similar patterns were observed for ancestors of *Acropora digitifera*, a coral, presumably owing to similar ecological dependencies. However, since the LGM, the coral population experienced growth while the anemone and symbiont experienced decline then stasis.

During a prolonged interval prior to domestication, potato ancestors decreased by half while the ancestors of the agent of potato blight doubled, suggesting they were independent prior to their joint population explosion since the advent of potato cultivation (Figure 1 and Supplemental Information). This result reinforces prior data demonstrating recurrent shifts among species of *Phytophthora* to unrelated plant hosts, and indicates that the association between potatoes and *P. infestans* in Central Mexico and the Andes was established late in the evolution of the fungal group, restricted to members of a poorly differentiated species complex (“clade 1c”22). Other, related fungal species infect evolutionarily and geographically disparate host plants (e.g. Malaysian epiphytes, European clover, North American Douglas Fir). Here, contrasting demographies prior to potato cultivation help date this historically consequential event in plant pathology, illustrating the power of comparative demography as a decision tool when considering the longevity of parasitic relationships.

## Discussion

Our approach builds on prior achievements extracting demographic information from heterozygosity patterns. Genomic regions whose evolutionary history has been discretized by subsequent recombination harbor distinctive amounts of heterozygosity, enabling inferences about the effective population size assuming a model of time-to-coalescence. In principle, this approach can be used to examine the degree of correspondence among any pair of high-quality genome assemblies in which heterozygous positions have been identified, provided that genome scaffolds are of sufficient length to capture the requisite recombination blocks. Here we reexamined published inferences derived from PSMC (in the cases of wild boars and primates), performed our own genome sequencing and analysis (in the case of *T. spiralis*), and mapped reads to existing assemblies that had not been previously subjected to PSMC analyses (anemone and its dinoflagellate symbiont, falciparum malaria, *Anopheles* mosquitos, potato, *P. infestans*, and some wild boar and domestic pigs).

When introducing PSMC by application to human genomes, Li and Durbin^6^ discounted the most recent 20,000 years of reconstructed history as unreliable, and most subsequent analyses have followed suit, leaving unexamined the ascendance of countless species (our domesticates, weeds, pests and pathogens) and the demise of countless others (by hunting, habitat destruction, pollution, and climate forcing) during the Anthropocene. We note, however, that while only ~800 human generations account for the last 20,000 years, 800 generations represent far less time for shorter-lived organisms. Reproducible signals of population growth, since the advent of agriculture, are clear in each domesticated host (and each of their pathogens) examined here, justifying an important extension of PSMC to the eventful recent past.

Obligate parasites cannot survive except in their hosts, providing a strong *a priori* basis for assuming shared demographic fates. However, even with strong and sustained correlation, alternative causal explanations merit consideration. For instance, climate variation engendering glacial advance and retreat certainly influenced the abundance and distribution of myriad species lacking parasitic or symbiotic dependencies; indeed, leaf-eating primates and pandas appear to have responded, in concert, to shared environmental imperatives^23^. While demographic parallels cannot prove any particular hypothesis as true, starkly contrasting demographic histories (such as those distinguishing potatoes from the fungal agent of potato blight, prior to potato domestication), can decidedly undermine the case for evolutionary stability in given parasitic relationships.

Our approach fruitfully addresses a longstanding question separating ecology from evolutionary biology: Do symbiotic and parasitic relationships long endure? Our demographic approach complements phylogenetic approaches, providing means to examine an interval lasting more than 100,000 years, beginning as early as 1 MYA and continuing as recently as 400 generations ago. Doing so affirmed parallel demographies in three case studies (malaria, *T. spiralis*, and a marine symbiosis) and identified a notable exception (substantiating and helping date the origins of potato blight). The rapidly expanding repository of high-quality genome assemblies afford newfound opportunities to consider the longevity of myriad parasitic and symbiotic relationships. Countless other biological dependencies (i.e. plants and their pollinators, predators and their prey, herbivores and their forage, and any pair or group of organisms engaging in synergistic or antagonistic relationships) may be similarly examined with these tools, vastly enriching our understanding of evolution, ecology, and epidemiology.

## Methods

Here, we describe in detail our means of acquiring sequence reads, assembling them into scaffolds, calling heterozygous positions, and analyzing the distribution of heterozygous positions using the pairwise sequentially Markovian coalescent model (PSMC, Li and Durbin 2011). We then describe methods for assessing the degree of correspondence between pairs of demographic plots, and determining the extent to which synchronized and natural pairs of plots better correspond than do plots of arbitrary species pairs or natural pairs lacking proper temporal order.

### Sequencing

Short-read whole genome sequencing was used to produce novel genome assemblies of *Trichinella spiralis* isolates from Asia and Europe. Isolates were collected from carcasses of animals determined to be infected post-mortem. Western samples were collected for molecular epidemiology purposes by the U.S. Department of Agriculture. Genomic DNAs from Asian isolates were graciously provided by Dr. Mingyuan Liu, Jilin University, Changchun, China. Genomic DNA was extracted using Promega DNA-IQ magnetic bead technology with the Tissue and Hair Extraction Kit Protocol according to manufacturer instructions. Extracted DNA was fragmented using Illumina’s NexteraXT DNA Library Prep Kit according to manufacturer instructions. After quantifying libraries, paired-end 300-base pair sequencing reads were generated on an Illumina MiSeq using the Illumina MiSeq Reagent Kit v3 chemistry. Each read was trimmed such that no base would have a higher than 5% chance of an error, and only two bases with quality scores less than Q20 would pass into the final read. Following quality control, reads longer than 50 bases were mapped to reference scaffolds using the Geneious Assembler with default settings in Geneious v.10.2.3.

### Previously published genomes

Genomes from species of interest were obtained from the public databases hosted by the National Center for Biotechnology Information at the National Institutes of Health. For each species, the chromosome or scaffold sequences were downloaded for use as reference sequences. Corresponding sequence read archives (SRA files) from individual samples were downloaded and subjected to quality control measures described above. Individual reads were mapped to the chromosome/scaffold sequences using Geneious Assembler in Geneious v.10.2.3. Accession numbers for scaffolds/chromosomes and sequence read archives (SRA files) are listed in Supplemental Table S2.

### Variant-calling and determination of consensus sequences

Variant-calling was performed on all assemblies using Geneious v10.2.3. Consensus base calls were derived from the 75% of reads with the highest representation at any particular locus. If sequencing depth at any base was less than 5X, that base was considered indeterminate (N). The consensus for each chromosome was exported in FASTA format for use in subsequent analyses using IUPAC nomenclature to denote heterozygotes.

### The Pairwise Sequentially Markovian Coalescent (PSMC)

Our approach examines whether, to what extent, and when pairs of PSMC plots correspond. The estimates of PSMC derive from the distribution of genomic blocks that are defined by a given level of heterozygosity. The underlying coalescent model considers the stochastic processes of mutational accumulation and the persistence, or loss of ancestral alleles. It further considers the effects of recombination in breaking down blocks of inheritance. Against the frequency distribution of heterozygous blocks expected for genomes descended from ancestral populations of constant size, actual distributions of heterozygosity are considered. Disproportionately frequent occurrence of blocks defined by a given level of heterozygosity is taken as evidence that the ancestral population waned; the date of such a bottleneck is estimated from the temporal interval over which that level of heterozygosity would be expected to accumulate. Similar intuition governs this model’s estimates of episodes of population growth (during which fewer-than-expected coalescence events would have occurred). The end result of any PSMC analysis is a plot of effective population size through time, limited by the detection of adequate numbers of coalescent events in the recent past, and the ability of the algorithm to identify recombination blocks from the remote past.

PSMC was run on each genome assembly individually using the default parameters. For final PSMC plots of effective population size through time, a minimum of 100 inferred coalescent events were linked to obtain robust size estimates in the geologically recent past, a threshold ten times more stringent than previously recommended to avoid overfitting.

The output of PSMC is broken into a specified number of “atomic time units,” each with associated parameters representing relative time and population size. We calculated the slope over “atomic time intervals” as the difference in N_e_ between adjacent atomic time units, divided by the difference in years, after scaling to a default annual mutation rate of 2.5e^−8^. Because atomic time units are initially scaled relative to the specific inferred 2N_0_ of a given sample, distinct samples (even replicates of a given species) are not initially synchronized. In order to compare contemporaneous slopes, we specified a set of standardized time-points: one for every thousand years between the present and 400,000 years ago, the last whole thousand-year atomic time point represented in the PSMC results of host samples under consideration based on the species with the shallowest history. Similar results were obtained for 135,000 years.Each standardized time-point was then assigned to its appropriate atomic time interval for further consideration of population size change, allowing multiple time-points to fall within the same atomic time interval (defined by adjacent atomic time units).

### Comparing demographic plots

In order to quantitatively assess how closely the demographic histories of two populations paralleled each other through time, we scaled PSMC results into absolute terms of years and effective population size (N_e_). This scaling requires specified generation time (g) and mutation rate per generation (µ). Uncertainty characterizes the value of these parameters for some of the organisms studied here, but given a biologically plausible range, the true value might be approximated by the one that minimizes differences between the PSMC curves, suggesting that they are appropriately aligned.

Previous work used the Pearson correlation coefficient (R) to evaluate the similarity of two PSMC curves. This approach is appropriate when proportional changes in population size are expected, *a priori.* However, contemporaneous changes in the size of a host’s and parasite’s populations might not occur in a strictly proportional way; there are typically many parasites or symbionts for each host, which drive large differences in their absolute rates of population growth and decline, and might induce non-linear responses. Indeed, in initial tests, we found Pearson’s R to be very sensitive to dramatic excursions in the PSMC plot, particularly (as was always seen for populations influenced by human activity since the inception of agriculture) populations that grew explosively in the recent past; biologically implausible parameter estimates could thereby result.

To address this problem, we derived a curve-comparison algorithm that minimized contemporaneous differences in slope sign between PSMC plots of two species or sets of organisms based on a simple ternary system of negative growth, stasis, or positive growth (assigned values of −1, 0, and 1, respectively). This procedure sought to maximize temporal intervals of shared population growth or decline, without regard to the magnitude of such growth or decline.

For each species pair considered, the host curve was held constant while the relative generation time was allowed to vary for each dependent species from 0.001 to 1.000 in 0.001 increments, assuming a constant generational mutation rate. This varied the temporal scaling of atomic time units without changing the shape of the curve. For each incremental change in generation time, the slope sign (−1, 0, or 1) at every 1000 year time interval was calculated. For example, if a species population contracted during the atomic time interval covering 10,600 to 7,300 years ago, then the time-points of 8,000, 9,000 and 10,000 years would receive a slope value of −1. Intervals where the absolute magnitude of slope was below the 10th percentile were assigned values of 0, signifying stasis. After determining slopes for all time-points, the difference in slope between putative species pairs was calculated for each time-point, ranging from 0 to 2. The average slope difference was recorded as a curve-fitting metric, where values of 0 indicate a perfect match and 2 an inverse correlation. We also recorded how many time-points the average slope difference calculation was based on and how many distinct atomic time units, from the PSMC output, those time-points intersected. The generation time that minimized slope difference while engaging at least 10 atomic time units was taken as the best curve fit between any two species if it fell within a plausible range (Supplemental Table 3). This approach slightly reduced noise and accounted for a degree of uncertainty in the PSMC estimates of effective population size at each point in time.

The correspondence between the demographic histories inferred for extant host- parasite relationships was compared, using this curve fit metric, to alternative pairs in the malaria and *T. spiralis* systems. For example, falciparum malaria now infects human beings, but its demographic history was also compared to that inferred for gorillas, considered the ancestral host for this parasite lineage. In the case of *T. spiralis*, parasite demography was considered in light of distinctions previously noted in the demographic histories of *Sus scrofa* in Europe and Asia (Figure S2). After completing the curve-fitting for all plausible hosts within a system (e.g. malaria with humans or gorillas), the host with the smallest average slope difference was deemed the best fit, and thus, the “natural host” responsible for controlling the growth or decline of the parasite of interest over the timeframe encompassed by the PSMC plots (Figure S3). This designation was used for further analysis below.

Finally, the relative generation times for each dependent species could then be translated into actual time units (years or months) by assuming appropriate mutation rates and generation times for the best-fit host. We note, with interest, that this same method could be used, instead, to estimate relative mutation rates for any pair of species for which generation times are known *a priori*.

### Testing curve fit

Any curve-fitting procedure might be capable of fixating on spurious parallelism, even among species pairs sharing no true history. We therefore sought to verify that observed parallels were more than chance occurrences.

One means to do so entailed determining whether a parasite’s population growth more closely mirrored contemporaneous growth in its host than it mirrored growth in its host at arbitrary times. For example, if the entire reconstructed history of a host was dominated by growth, every time-point would have the same slope and the order would be irrelevant. To carry out this test, we randomized the slopes of each sample with respect to time, separately, for each set of samples. Correctly ordered slopes from one species were then compared to non-chronological ones from the other. A two-tailed t-test was used to evaluate whether the mean slope differential for contemporaneous plots were less than when one plot’s temporal order was randomized with respect to the other, based on three independent randomizations (Figure S1). In general, the more complex the curve (i.e. the more local maxima/minima) the more power we should have to determine goodness of fit.

In another attempt to evaluate the extent to which shared histories likely explained parallel demographic plots, we compared the fit of natural species pairs (defined by extant parasitic or symbiotic relationships) to arbitrary species pairs (Figure S2). For example, there would be no strictly biological reason to expect parallel demographic histories for *P. falciparum* with pigs, *T. spiralis* with potato, *P. infestans* with gorillas, or this species of *Symbiodinium* (*S. minutum*) with a species of coral (*Acropora digitifera*) with which it has not been shown to establish a symbiosis. In so doing, we sought to test whether our curve-fitting approach was so flexible as to identify parallels where none should have been expected, *a priori*. Thus, each parasite/symbiont PSMC plot was fit sequentially to all host species in the entire study as described above. At the optimal fit for these implausible hosts, the average slope difference was calculated (Figure S2). The average slope difference of parasites to all natural hosts, other plausible hosts, and implausible hosts was graphed with 95% confidence intervals combining all four systems (Figure S3).

### Code availability

Custom scripts were written in Python and R for evaluating fit of curves. These will be made available to reviewers upon request and placed in a public repository (GitHub) following acceptance of the manuscript.

## Supplemental Information

This supplement provides additional details about the samples, subsidiary analyses, and original research results that explore and expand on the capabilities and limitations of comparative historical demography as elaborated in the associated manuscript. We detail more specific findings related to the four biological systems studied here (malaria*, Trichinella spiralis*, *Symbiodinium minutum*, and potato blight systems), emphasizing themes such as replicability, specificity of a parasite’s fit to a particular host species, geographic specificity of host-parasite demographic parallels, and the applicability of PSMC to organisms (such as potatoes and most other crops) where successive genome duplications and uncertain genome assemblies may inflate estimates of heterozygosity and thereby bias estimates of ancient population size.

### Effect of variant-calling methodology on PSMC

We tested the sensitivity of PSMC to alternative variant-calling methods, which may influence the number and distribution of heterozygotic base calls in the consensus sequence. We observed little difference in shape or scaling between PSMC plots of *T. spiralis* isolates based on heterozygote-calling performed, alternatively, by Geneious or the mpileup function of Samtools (samtools mpileup -uf Genome.fasta test.sorted.bam | bcftools view -cg - | vcfutils.pl vcf2fq > consensus.fq) (Figure S4).

The heterozygote frequencies found in the PSMC input files (.psmcfa) generated from these consensus fasta files were also similar irrespective of the variant-calling program, with the Geneious consensuses generally having a greater proportion of missing data, suggesting the filtering thresholds we applied (75% consensus, >5x coverage) were slightly more stringent than the Samtools defaults (Supplemental Table 4). Nadachowska-Brzyska et al. (2016)1 reported that increasing their per-site filtering thresholds shifted their PSMC curve toward the present, and this effect is also apparent to a minor degree in our plots. The same authors found that more stringent filtering helped to resolve population fluctuations in PSMC plots, though the significance of this faded for average read depths greater than ~18x.

In general, joint histories of species pairs could be traced from about 500,000 years ago (where years result from joint estimates of mutation rate and generation time) until a few thousand years ago. In considering the evolutionary longevity of particular parasitic or symbiotic relationships, our methods appear well-suited to that interval (and no longer). It was often the case that one, but not both, genomes in a given pair provided temporally deeper inferences on demographic history.

## Malaria system

The origin and expansion of the agent of malaria, *P. falciparum*, is tied to human demographic history^2,3^. The most recent evidence indicates that *P. falciparum* shifted from gorillas to humans between 60,000 and 130,000 years ago^4^, though chimpanzees were previously suspected as the proximate source^5,6^. When humans began clearing land for agriculture, anopheline mosquito numbers increased accordingly, resulting in increased malaria transmission and presumably increased *P. falciparum* population sizes. Therefore, we sought to examine the demographic changes written in the genomes of hosts, vectors, and parasites to confirm our understanding of falciparum malaria in humans.

*Plasmodium falciparum* is only briefly diploid in its anopheline mosquito definitive hosts, and to our knowledge has not been sequenced in that life-history stage. Therefore, we mapped the distribution of potentially heterozygous positions by creating “pseudodiploid” genomes in silico by mapping short reads from two haploid sequencing projects derived from *P. falciparum* isolated in the same geographic region to the *P. falciparum* 3D7 chromosomes (the most complete representation of the *P. falciparum* genome to date). We took these assemblies to represent the joining of haploid gametes into transient diploid ookinetes. We selected sequencing projects with approximately equivalent numbers of reads in order that heterozygous bases could be determined based on read depths as above, assuming comparable coverage of chromosomes by each deeply sequenced genome. The resulting consensus chromosomes were used for downstream analysis, including estimates of relative generation time (Figure S5) that well match empirical estimates based on clinical and entomological observation. Producing pseudodiploids in this way, we note, could render any haploid organisms amenable to PSMC analysis.

Plasmodium falciparum notably parallels the bottleneck evident for human beings and their subsequent ascendency in the last ten thousand years; by comparison, other great apes have experienced stable or declining effective population sizes since the Neolithic (Figure S6). Similar demographic patterns have been inferred for primates circa 100 KYA-1 MYA, and would be expected to eventually converge on the shared demography of our common ancestors; subsequent to divergence, it would not be surprising to find that various primate species were subjected to common beneficial and injurious environmental conditions.

Chimpanzees had previously been considered the penultimate host for the ancestors of falciparum malaria until more closely related parasites were discovered in gorillas. Because ancestors of chimpanzees may have served as earlier hosts to ancestors to falciparum malaria, we considered the demographic history of *P. falciparum* in relation to that published by Prado-Martinez et al. (2013)^8^ for various populations of chimpanzees (*P. troglodytes ellioti*, *P.t. schweinfurthii*, *P.t. troglodytes*, *P.t. verus*). While all Old World primates (including *H. sapiens*) share certain features of demographic history, the histories of various sub-populations of chimpanzees notably differ (Figure S6). The demographic history of *P. falciparum* tracks human demography best, especially during the most recent historical interval, when human populations grew exponentially while other great apes did not. However, of the considered chimpanzee populations, *P.t. schweinfurthii* was the most consistent with human and malaria demography in the range of 80K – 1M years ago.

### Trichinella spiralis system

*Trichinella spiralis* is contracted by the consumption of meat in which larval parasites have encysted. Adult pairs mate within weeks, and disseminate new larvae to the tissues whereupon the next generation of larvae establish chronic infections that endure for the life of the host. Thus, infections are acquired after weaning and are transmitted at death. The mean generation time of the parasite would be shortened to the extent that infection reduced swine longevity and to the extent that the parasite also exploits shorter-lived hosts, such as rats. The curve-fitting procedures described above suggested ~ 4 parasite generations per wild boar generation (or 1.1 years, assuming 5-year generations for wild boar).

The inferred relative generation time of *T. spiralis* was inconsistent in Asia and in Europe (Figure S7), probably owing to limited complexity and depth in the demographic history reconstructions of European *T. spiralis*. Since domestication, pigs have undergone a population explosion, and our analysis illustrate commensurate exponential growth in the size of the parasite population (Figure S8). Wild boar have interbred with domesticated pigs, leaving tracts of homozygosity in wild boar genomes^9^ that may inflate estimates of recent population growth in wild boar.

As stated, the demographic plots for western *T. spiralis* were generally lacking in complexity, depriving curve-fitting procedures of information needed to achieve precise estimates (Figure S7). Nonetheless, their optimal fit is consistent with a decline in pig populations in both Europe and Asia that has previously been interpreted as having been caused by the LGM^10^. The inferred histories of Asian *T. spiralis* reach deeper in time and show greater demographic variability over time. Most notably, in contrast to European *T. spiralis* and to both European and Asian *S. scrofa*, the effective population size of Asian *T. spiralis* does not decline over the last ~50,000 years, but stabilizes and grows. All pigs and *T. spiralis* eventually gave way to dramatic growth over the Holocene.

### Symbiodinium system

A long-term demographic association between the anemone *Aiptasia pallida* and its symbiont *Symbiodinium minutum* might not have been expected owing to expulsion and die-offs of symbionts during periods of environmental stress^11^. Cnidarians, including anemones and corals, sometimes acquire new species after heat-induced bleaching^11^. Although such associations might therefore prove ephemeral, the population of *S. minimum* tracked that of *A. pallida* remarkably closely (Figure S10). The demographic histories of all three species were similar until approximately 10-20 KYA, following the LGM, when the coral population began a period of extreme growth while the anemone and symbiont populations began to decline.

We employed equivalent curve-fitting procedures so as to determine the best-possible demographic match over time, and derived a relative generation time of ~0.5 against the actual host, *Aiptasia pallida*, and 0.97 against the apocryphal host, *Acropora digitifera* (Figure S10). Notably, the estimated slope difference between host and symbiont demographic curves was less for the natural pair at almost every considered value of relative generation time (Figure S9); the average slope difference between *S. minutum* and that of its actual host (~.09) was notably less than to that of its apocryphal host (~0.17) when optimally scaled to each. Figure S10 depicts the histories of all three species, assuming a relative generation time of 0.52 for *S. minutum* (fit to *Aiptasia pallida*).

Their shared history in the Red Sea may partially explain the evidently shared demographic histories of the anemone and its symbiont (in contrast to that of the coral, obtained from Okinawa in the East China Sea). Although oceanic coral systems generally prosper when the global climate cools, glacier formation during the last glacial maximum resulted in a 120m drop in sea level, halving the surface area of the Red Sea, significantly reducing exchange with the Indian Ocean through the Strait of Bab el Mandab, and increasing the sea’s salinity by about 50%^12,13^. These events would likely have reduced the census size of the anemone population, as well as fragmenting it within the Red Sea and between the sea and ocean, further reducing the effective population size of the anemone. Smaller, fragmented host populations would likely have constrained growth of the symbiont. Subsequent rising seas may have engendered population growth of the anemone and its algal symbiont owing to increased habitat availability and increased gene flow among once-separated populations (Figure S10).

Considering the foundational role of corals in creating habitat for anemones and their symbionts, it is not surprising that the demographic histories of all three species are very similar in overall shape. Indeed, the histories of *Aiptasia pallida* and *Acropora digitifera* can be nearly superimposed by assuming a generation time ratio of 1:2 (data not shown). Although we have assumed a default generation time of one year for *Acropora digitifera* and *Aiptasia pallida*, for convenience and given a lack of information, *Acropora digitifera*’s generation time could be as long as its age of maturity (3-8 years^14^), which would imply a similar generation time for *S. minutum* and one of 6-16 years for *Aiptasia pallida*. That would be an implausibly long generation time for a dinoflagellate, but this relative scaling of demographic history could also be accounted for by a slower per-generation mutation rate. Indeed, the typical generational mutation rate of other haploid eukaryotic phytoplankton (~5e^−10^) is around 50 times slower than the mammalian one we have assumed by default (2.5e^−8^)^15^. This would imply an *S. minutum* generation time of approximately 3-8 weeks, rather than 3-8 years.

## Potato system

Infections of potatoes (*Solanum tuberosum*) with the potato blight *Phytophthora infestans* caused one of the greatest human disasters in agricultural history, resulting in the death or emigration of 20% of the Irish in the mid-1800’s. Potatoes originated in the Andes and may have been domesticated twice. Biogeographic data suggest the central Mexican highlands as the ancestral home of *P. infestans,* achieving global reach only after the potato domestication in the Neolithic and the globalization of agriculture beginning 500 years ago. *P. infestans* appears restricted to tuber-forming species of Solanum and does not easily cross with other species of *Phytophthora*. The closest known relatives to *P. infestans* (*P. andina, P. ipomoea, P. mirabilis,* and *P. phaseoli*) are also pathogens of plants native to the Neotropical highlands. However, other closely related congeners (*P. iranica, P. clandestina, P. tentacultate*) afflict diverse plant types (eggplant, clover, chrysanthemum) in diverse regions (Asia, Europe); an even greater diversity of host types and biogeographic regions characterize more inclusive groups of *Phytophthora,* including species infecting orchids native to Indonesia, lilacs in the Balkans, rhododendron native to the Himalayas, raspberry native to Northern Europe and Asia, and Douglas fir in Western North America. This broader evidence for host switching provides valuable context to our results, suggesting parallel demography only during the most recent past.

Prior to potato domestication and cultivation, potatoes and the agent of potato blight shared no obvious episodes of concerted population growth or contraction. Lacking such compelling information, and with a relatively short and simple reconstructed history of *P. infestans* to overlap, our curve-fitting algorithm easily identified a clear preferred relative generation time for the two at ~0.14 *P. infestans* generations per potato generation (which was assumed to be one year) (Figure S11). Adopting this temporal scalar resulted in an average slope difference (0.38 on a scale of 0-2) that was as poor as the best fit between apocryphal species pairs (such as wild boar), even with relatively tight constraints on plausible generation times for the pathogen (<3 months^16^). Among poor alternatives, the best-fit curves register by reference only to the recent (post-domestication) interval of population growth in potatoes. This is preceded by ~20,000 years of decline in potatoes and relative stasis in *P. infestans* (Figure S12). These findings are consistent with a close affinity of *P. infestans* with potatoes only since their domestication.

Noting an ancient tripling of chromosomal content in dicot plants^17^, we were careful to assess whether, how, and to what extent demographic plots might be biased if reads derived from paralogs in the potato genome were mismapped to the same position, thereby inflating estimates of heterozygosity. We examined this potential source of bias by filtering regions of the potato genome sequenced to especially high coverage (arguably over-represented owing to their occurrence as multiple copies). Prior to filtering, the potato genome dataset had an average sequencing depth of 31.8 reads per site with a large standard deviation (148.5). After filtering, the mean sequencing depth and standard deviation was reduced to 12.4 and 13.3, respectively. Filtering resulted in removal of approximately 1/5 of each scaffold for consideration by PSMC. After filtering, about 40 million bases of contiguous sequence in 10 scaffolds were available for analysis. Filtering high coverage areas reduced the overall heterozygosity in the potato genome from 1.19% to 0.895%. It also reduced the length of the scaffolds used for secondary PSMC analysis from 43,353,824 bp to 36,083,464 bp (17% removed from consideration). Confirming simulations entertained by Li and Durbin (2011)^18^, removing suspected paralogous regions did not affect the overall shape of curve, but did reduce the estimate of the most ancient populations (Figure S13). The essential demographic signal to remain stable, further substantiating the conclusion that potatoes and the fungal agent of potato blight did not experience parallel demographic histories until the age of potato domestication, beginning ~9,000 years ago.

---

**Figure S1:**
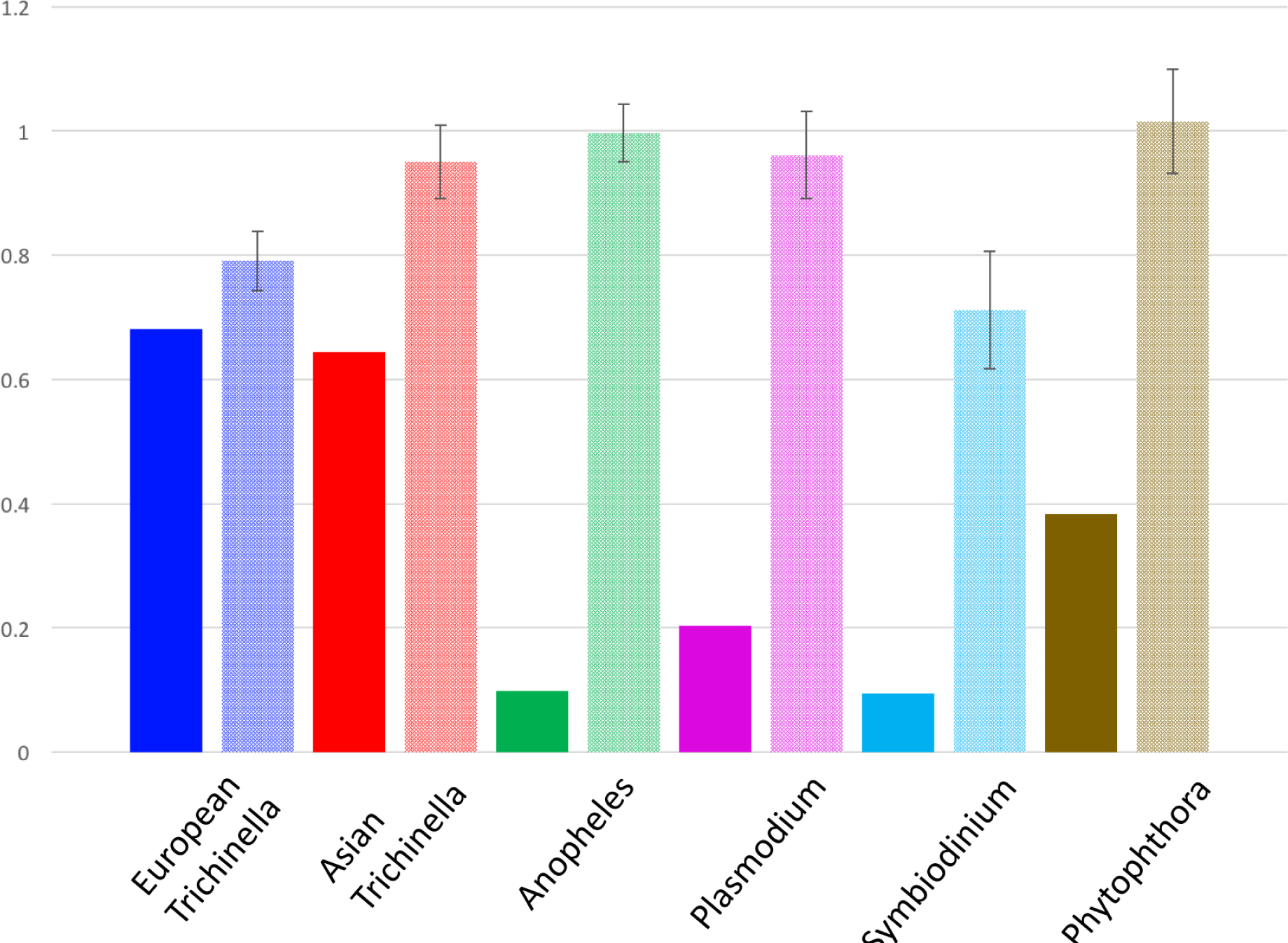
Comparisons of average slope difference between unmodified curves (solid color) and curves that have been randomized with respect to time (faded). Parasite or symbiont plots were aligned to their purported host plots (European *T. spiralis* vs. European *S. scrofa;* Asian *T. spiralis* vs. Asian *S. scrofa*; *P. falciparum* vs. human; *A. gambiae* vs. human; *S. minutum* vs *Aiptasia pallida*; *P. infestans* vs potato) and the average difference in slope direction was calculated. Subsequently, population size estimates of each parasite or symbiont were randomized with respect to time, and average slope difference and its variance were calculated with respect to the host. The 99.9% confidence interval is shown in association with the randomized curves, based on the variance among randomization replicates (n=3). Randomizing slopes with respect to time significantly worsened the match for every ecologically relevant pairing of species. Overall, randomization increased average slope difference by an average of 158% (p=0.0009).

**Figure S2:**
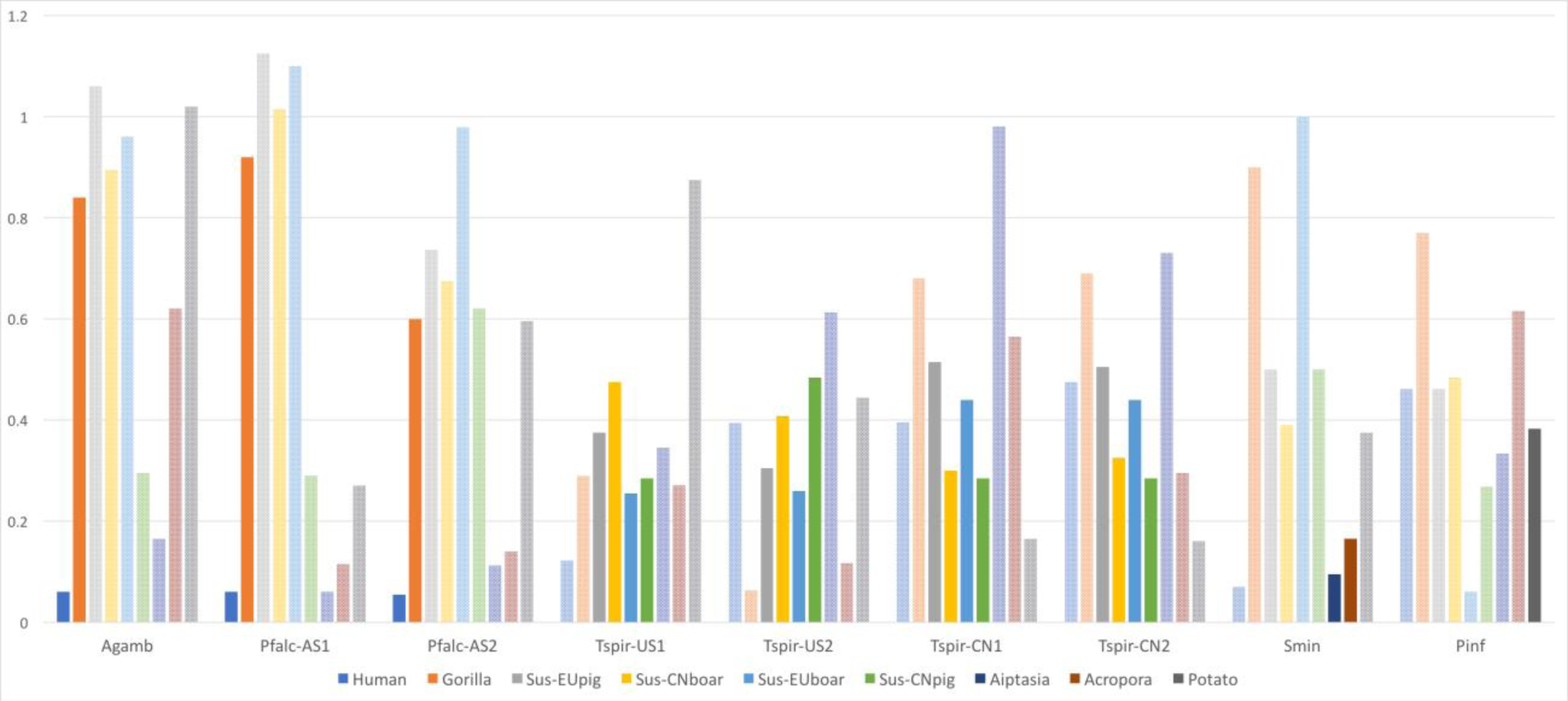
Comparison of minimum average slope difference for all considered host-symbiont pairs. Bars representing the fit of biologically plausible hosts are highlighted, while apocryphal matches are faded, but shown for context. In all cases, the true host is preferred from the arena of plausible hosts (e.g. European boar > Asian boar for European *T. spiralis*), but in some cases an apocryphal match scores better (e.g. European boar > potato for *P. infestans*).

**Figure S3:**
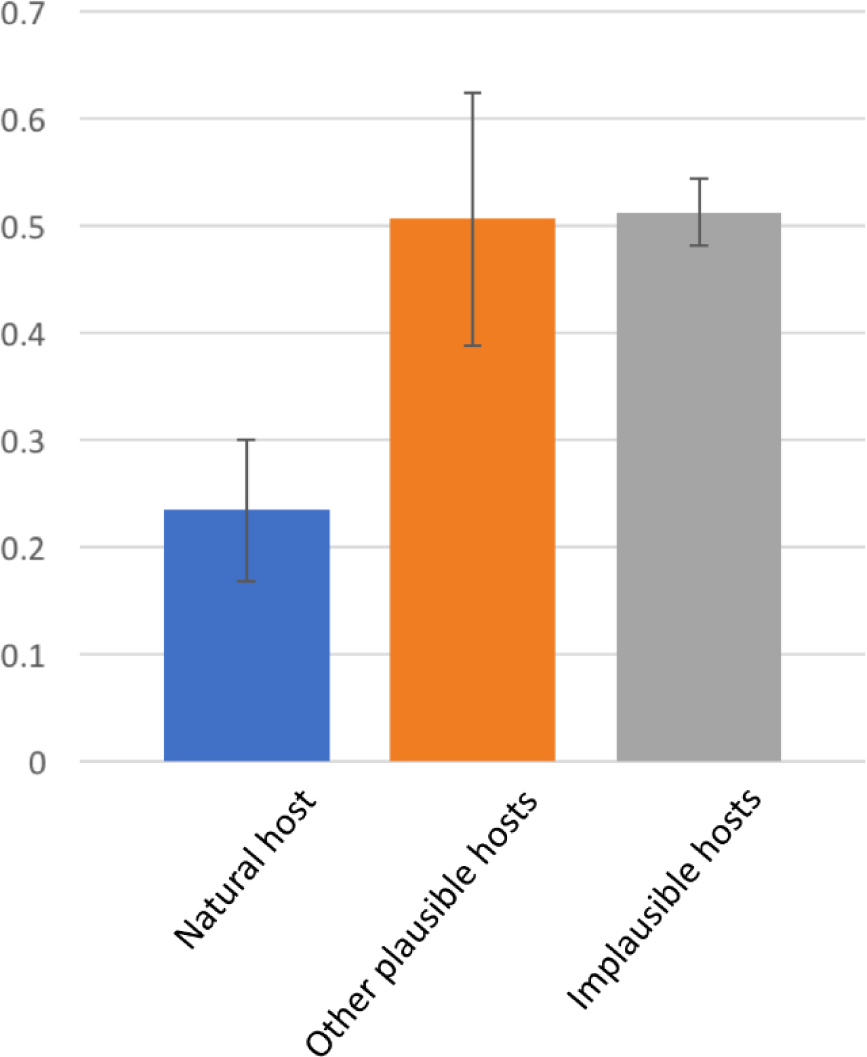
Comparison of minimum average slope difference for the natural host (e.g. *P. falciparum* x human; n=13) versus plausible (e.g. *P. falciparum* x gorilla; n=12) and implausible (e.g. *S. minutum* x human; n=56) alternative host-symbiont combinations. Error bars represent the 95% confidence interval of all comparisons. Overall, natural pairs exhibited around half the average slope difference of other pairings, while no difference was observed between plausible and implausible alternative pairings.

**Figure S4:**
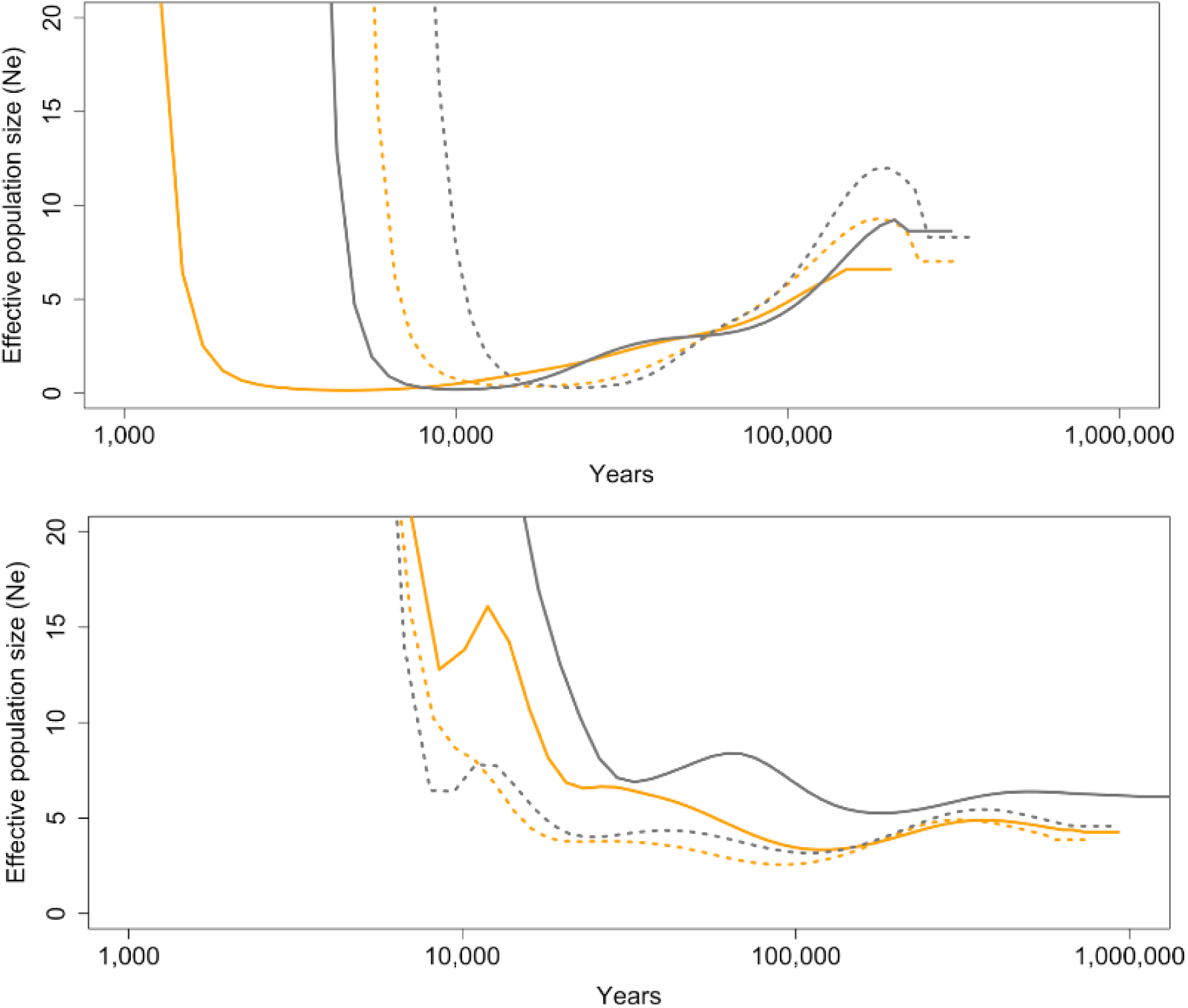
PSMC plots based on consensus sequences generated by Geneious (orange) and Samtools (grey), for two *T. spiralis* isolates from America (left) and two from China (right)

**Figure S5:**
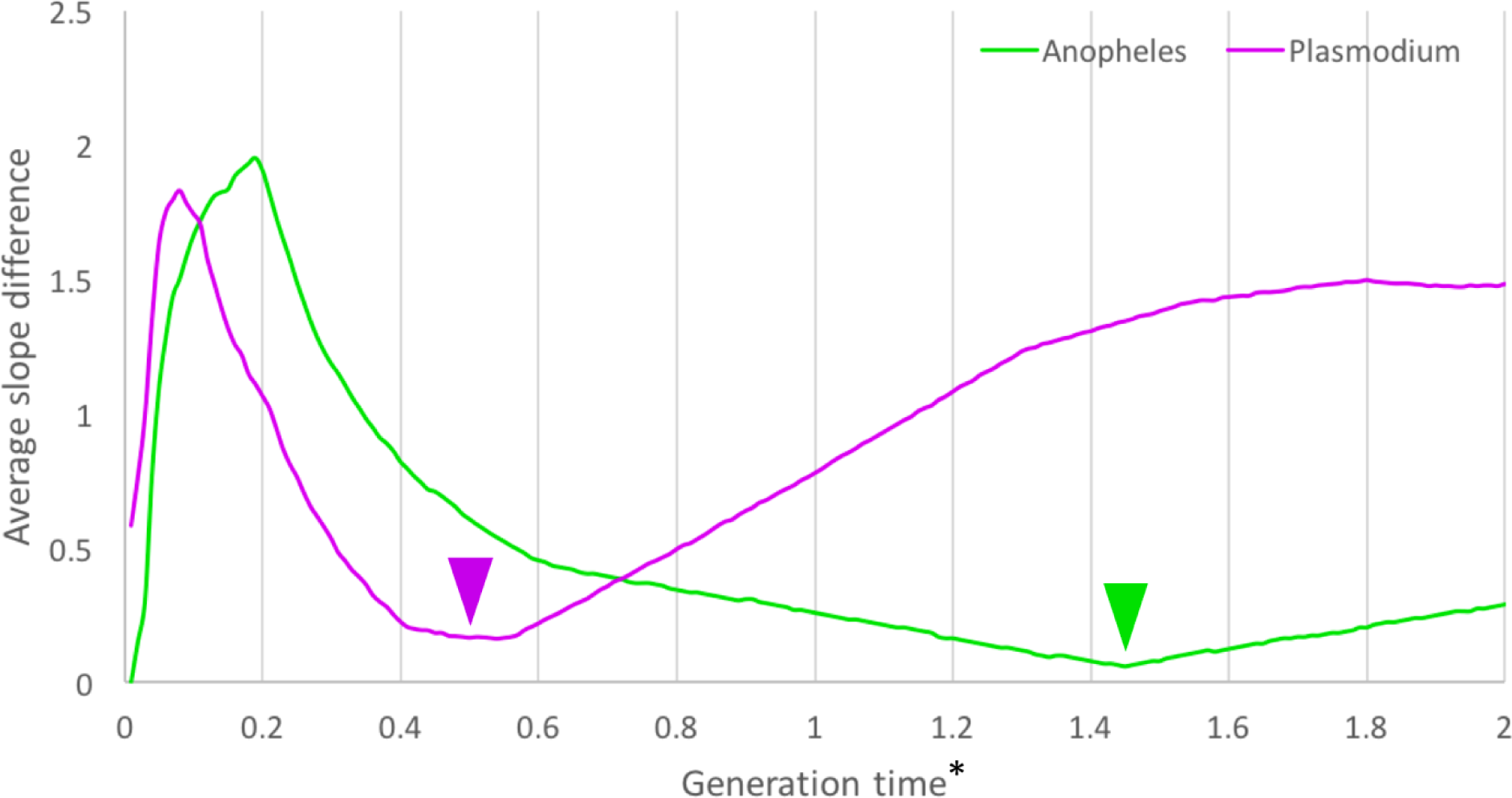
PSMC curve-fit to French Human as a function of generation time for *A.gambiae* (green) and *P. falciparum* (purple), assuming a generation time of 25 years for humans and identical per-generation mutation rates of 2.5e-8. The lowest average slope difference was found at 1.43 years/generation for *A.gambiae* and 0.48 years/generation (~23 weeks/generation) for *P. falciparum*. However, the slower *Drosophila* mutation rate (1.1e-9) may be more appropriate for mosquitoes7, which would imply an *A. gambiae* generation time of 0.057 years, or approximately one generation every 3 weeks, in line with empirical estimates based on knowledge of mosquito ecology.

**Figure S6:**
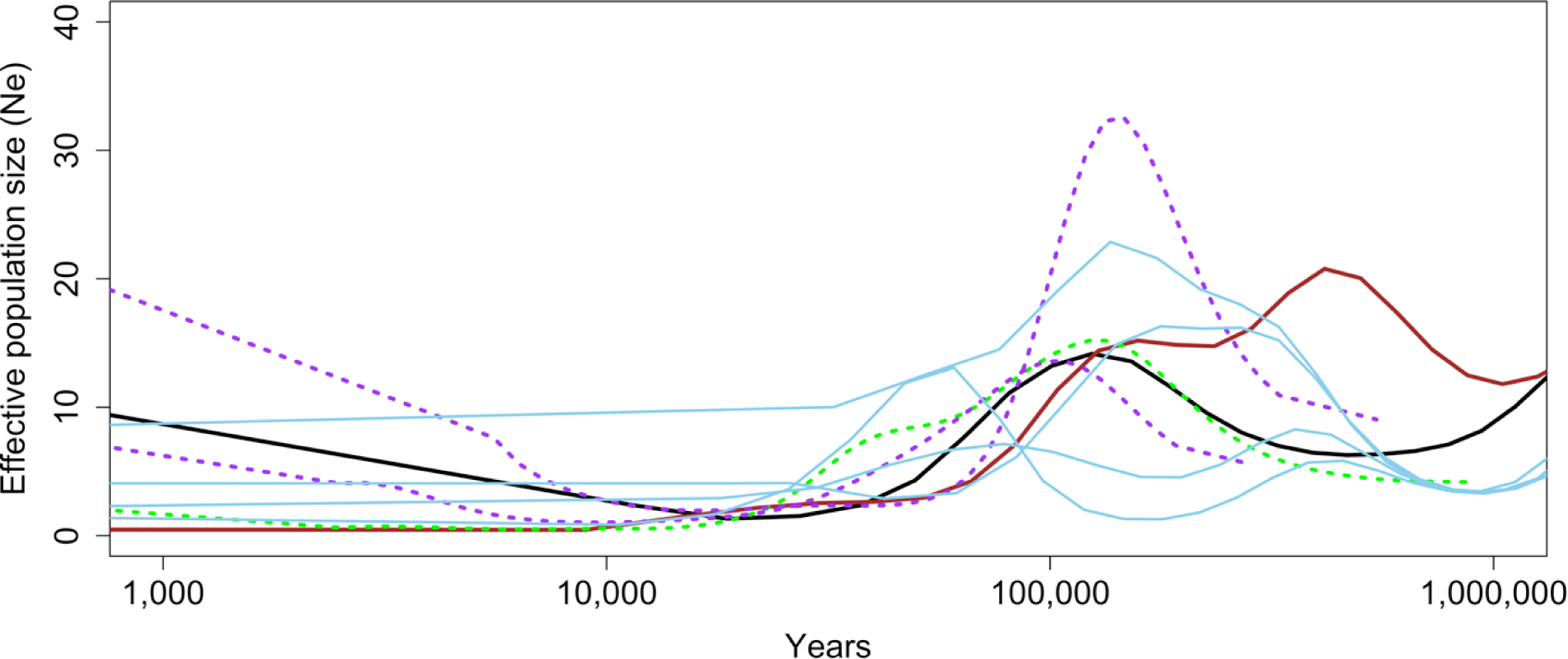
Estimated demographic histories for *P. falciparum falciparum* (violet) and *A. gambiae gambiae* (green) in relation to that for human (black) and gorilla (brown) and multiple subspecies of chimpanzee (blue). Gorilla populations grew and shrank in parallel with humans, Anophelines, and *P. falciparum*, with steep declines from 200,000 ybp to 70,000 ybp, followed by a small stable population between 70,000 and 11,000 ybp. From ~100-11 KYA, gorilla populations declined in concert; prior to then, their populations were notably larger and since then notably smaller. *P. falciparum* and mosquito closely parallel human demography for a considerable period, including growth in the most recent past unique, among primates, to human populations. Recent growth in the mosquito population, while present, is much less dramatic than that observed in the human and parasite populations. Note that human and gorilla effective population sizes were uniformly scaled up by a factor of 10 to be more easily visible in this figure.

**Figure S7.**
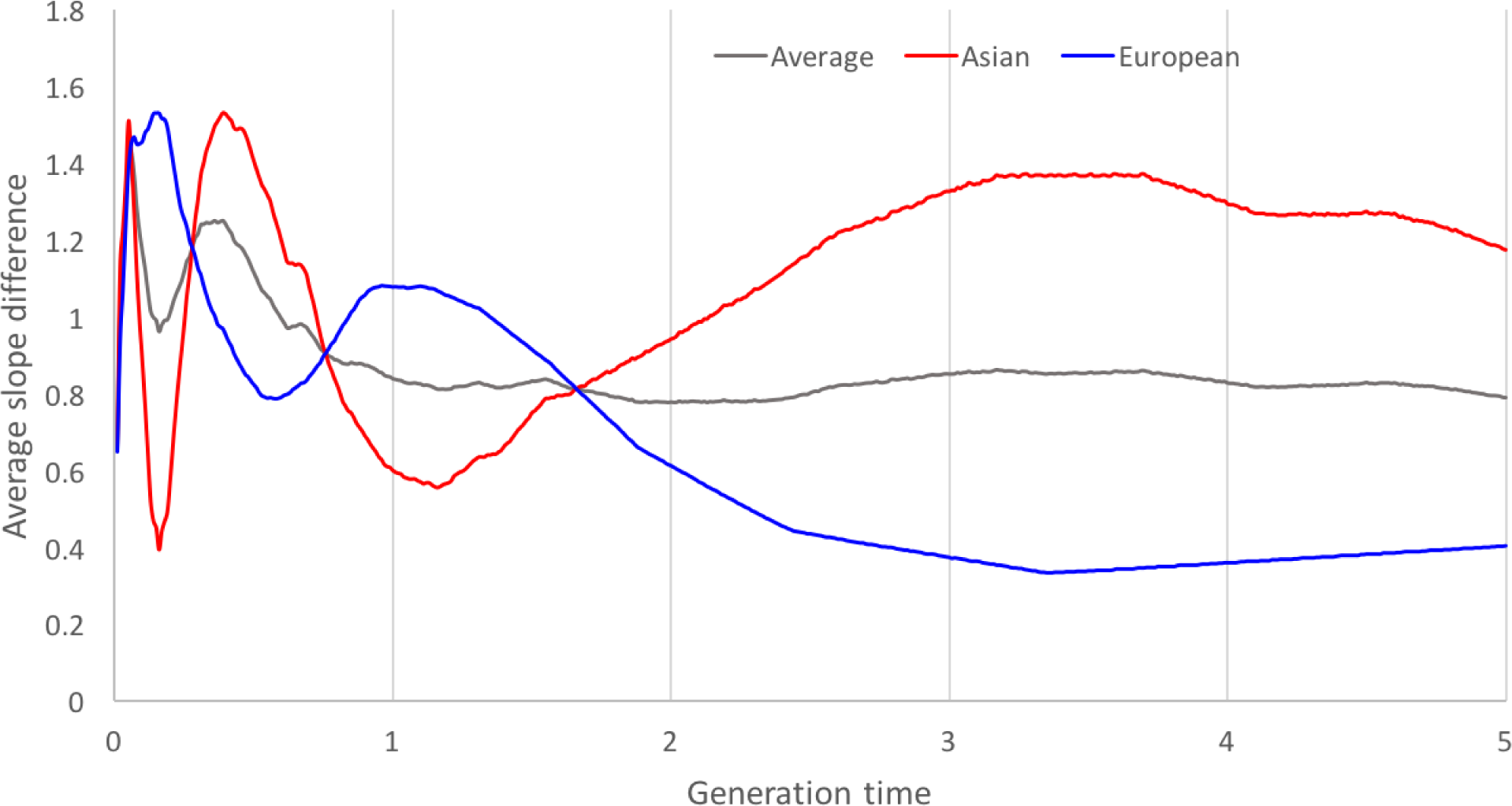
Fit of European (blue) and Chinese (red) *T. spiralis* to sympatric *S. scrofa* as a function of their generation time. Given 5 years per boar generation, the lowest average slope difference between the average of the two regional fits (grey line) stabilizes at approximately 1.1 years per *T. spiralis* generation, which is the regional optimum for the Asian *T. spiralis*. Because the Asian genomes record a deeper and more complex demographic history than the European ones, we elected to proceed with 1.1 years/generation as the species optimum generation time for *T. spiralis*. Note that although the Asian *T. spiralis* slope difference is minimized at <6 weeks, this biologically implausible result derives from few overlapping time intervals (between parasite and host).

**Figure S8:**
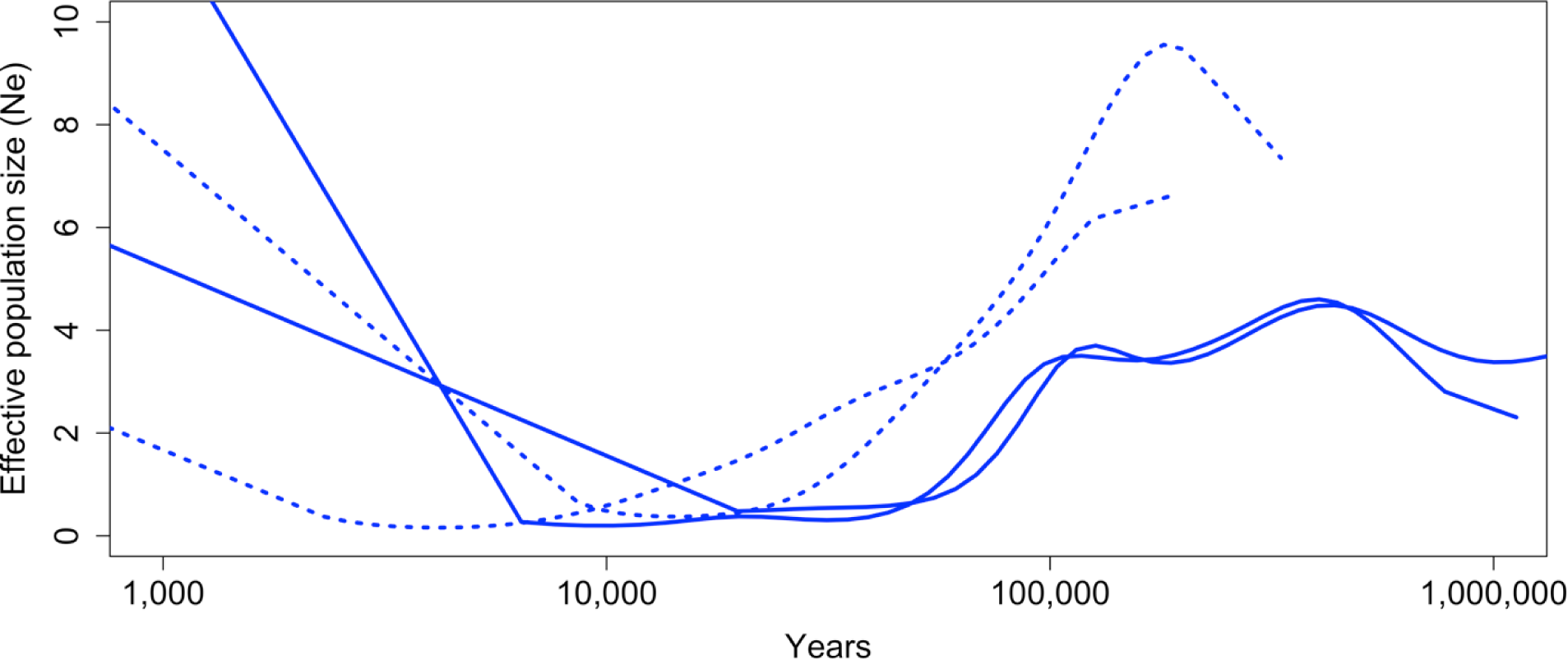
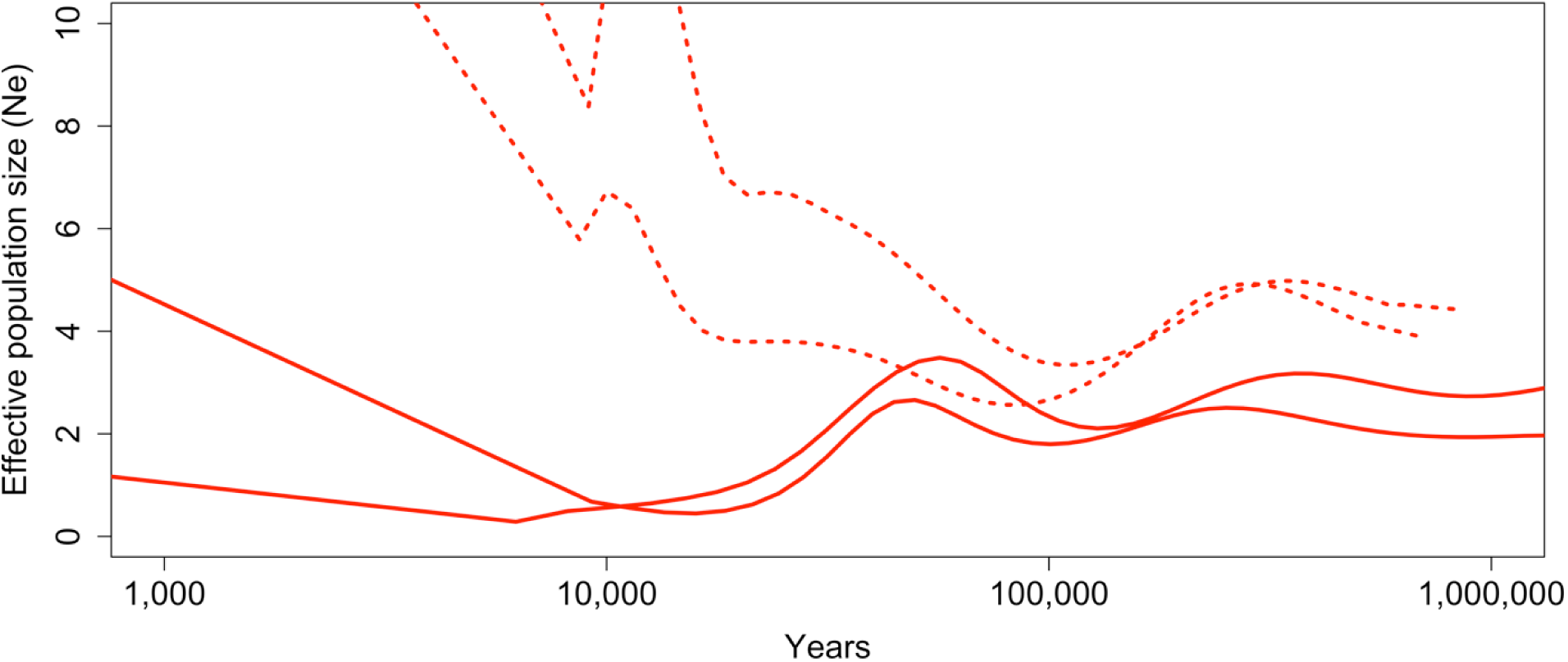
Regionally-specific reconstruction of *T. spiralis* demography in relation to sympatric wild boar. Regional distinctions in the histories of wild boar are recapitulated in the histories of the parasite prior to the age of swine domestication. Since domestication, parasite populations in each region have undergone precipitous growth. The PSMC curves are those reconstructed for wild boar by Groenen et al, 2012)^10^.

**Figure S9:**
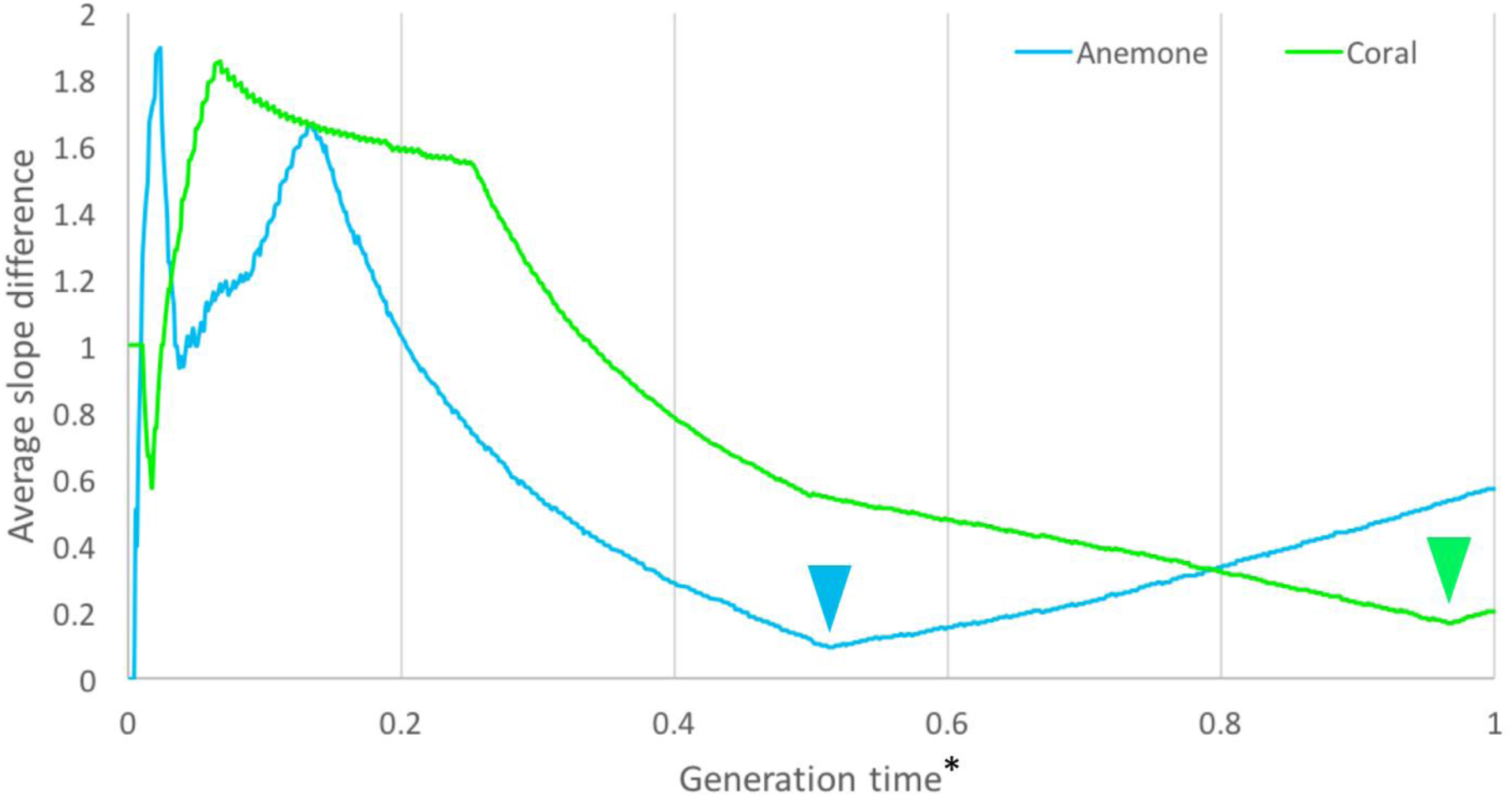
The slope difference between *S. minutum* and its natural host (*Aiptasia pallida;* blue line) was less than to another marine invertebrate, the coral *Acropora digitifera* (green line). The fit to *Aiptasia pallida* was optimized at a relative generation time of approximately half (0.52x) that of the host. Optimal fitting to *Acropora digitifera* would require almost identical generation times for host and symbiont (0.97x).

**Fig S10:**
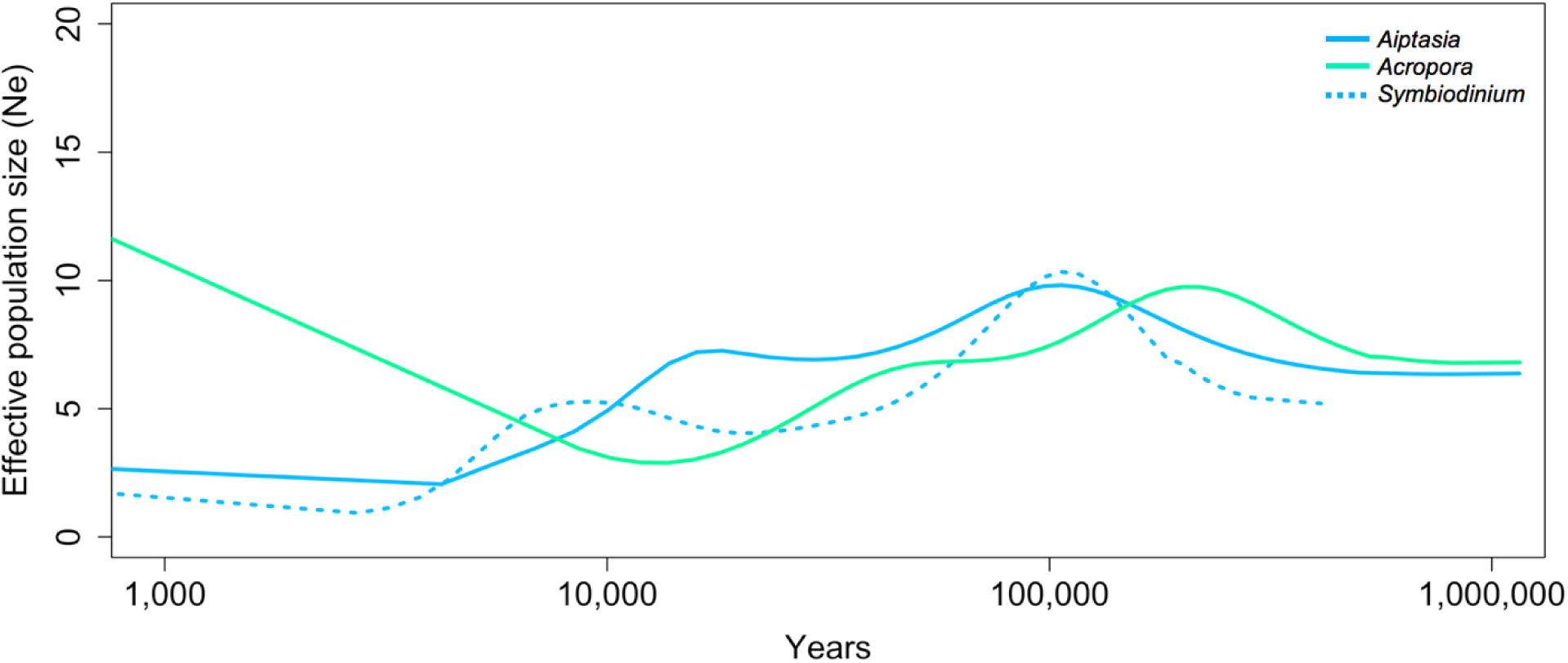
Demographic histories of coral (*Acropora digitifera*), anemone (*Aiptasia pallida*), and the anemone’s symbiont (*S. minutum*), assuming annual generations for the coral and anemone and biannual generations for the symbiont. Rises and falls of *S. minutum* (dotted blue line) approximate those for its extant host, *Aiptasia pallida* (solid blue line) for approximately 1 million years. A similar pattern of growth and decline is observed, albeit out of sync, for *Acropora digitifera*, but the demographic trajectory of *Acropora digitifera* separates in the last 20,000 years, experiencing rapid population growth while the anemone and its symbiont decline and stabilize.

**Figure S11:**
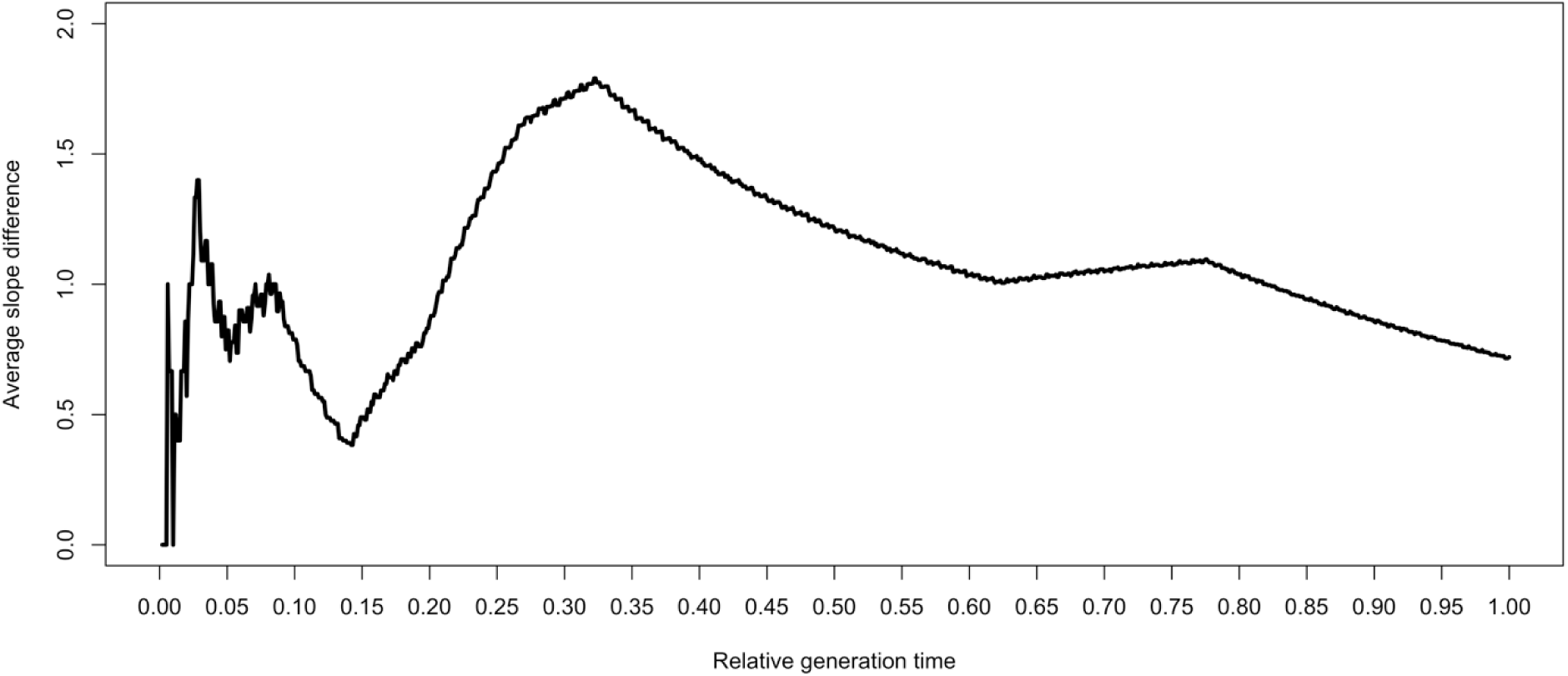
Fit of *P. infestans* demographic history to that of potatoes as a function of relative generation time. The optimum is reached when a single potato generation equals ~10 generations of *P. infestans.* This short generation time is based on concerted growth in the host and pathogen only since the domestication of potatoes in the last ~8,000 years (Figure S12), suggesting that shared demography may have been limited to this recent temporal interval.

**Figure S12:**
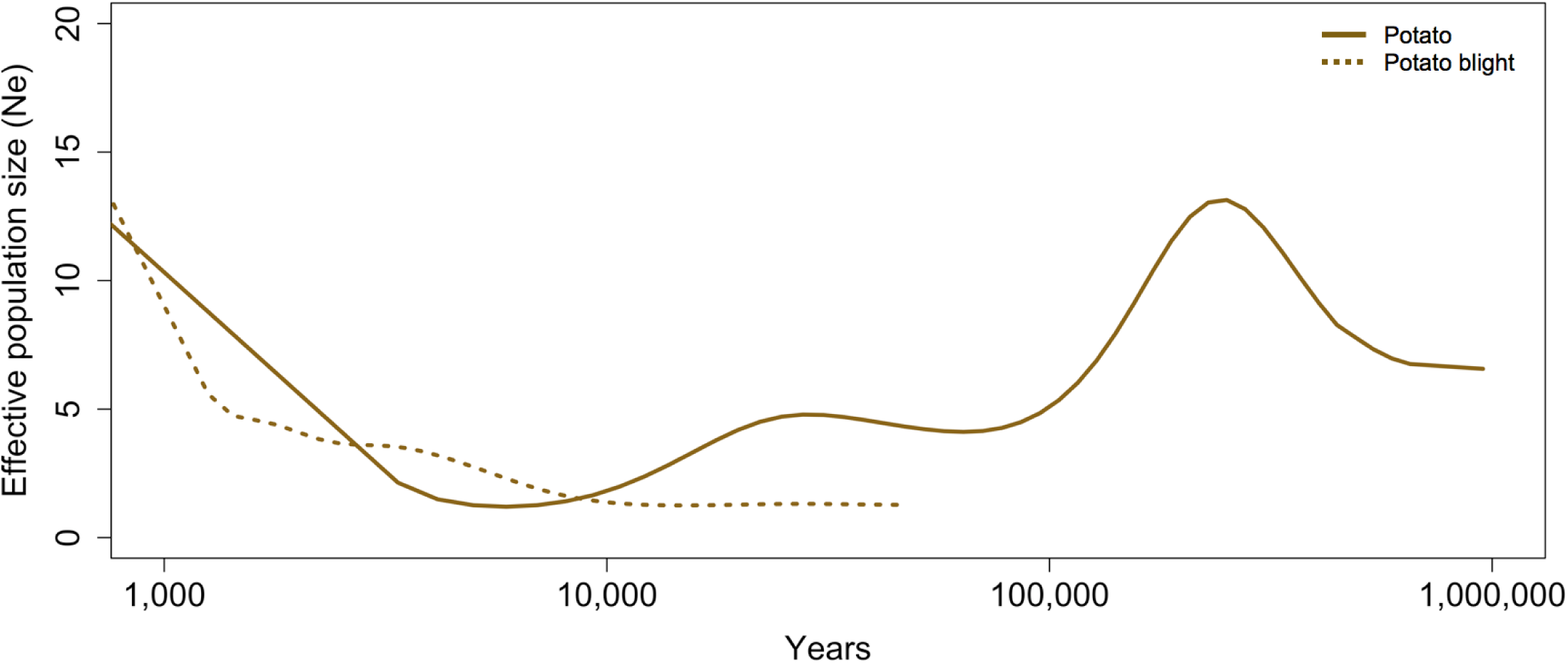
Contrasting demographic histories of potato and *P. infestans* the agent of potato blight excepting since domestication. The only demographic feature indicating shared history in this plant/pathogen system relates to recent growth in each. Here, the demographic history of potato (solid line) is depicted against the history of the fungal pathogen assuming the optimal relative generation time, which does not convincingly match the history of potatoes prior to potato domestication.

**Figure S13:**
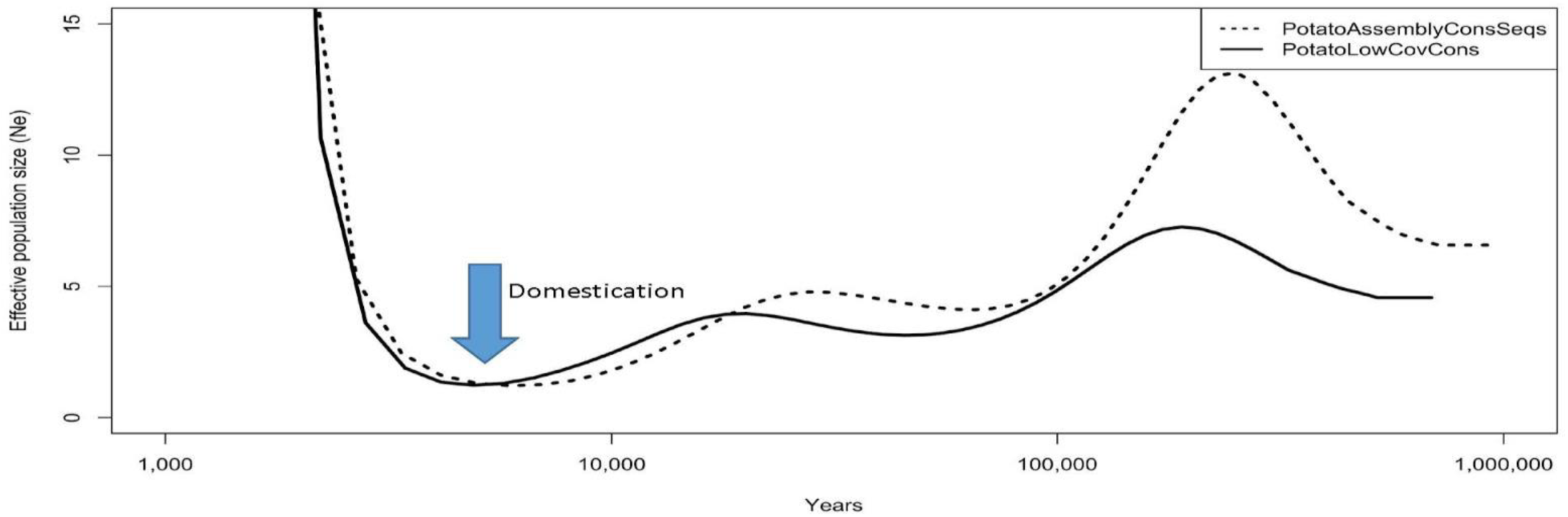
Effect of removing suspected paralogs from the potato genome prior to estimating its demographic history using PSMC. The PSMC plot derived from the potato genome assembly (dotted line) was compared to an equivalent analysis performed on a dataset from which anomalously deeply sequenced regions were removed (solid line), owing to the suspicion that paralogous gene copies derived from genome duplications might be inflating estimates of heterozygosity. As predicted (Li and Durbin, 2011)^18^, removing these sequences did reduce estimates of ancient population size. However, this quality control step did not fundamentally alter the inference as to when ancestral potato populations grew or contracted, and specifically had no appreciable effect on demographic estimates within the last 30,000 years.

**Table S1:** Host and symbiont species, sample abbreviations, and the form in which the sample was obtained. The *P. falciparum* isolates marked with asterisks were assembled as pseudodiploids from multiple sets of published reads, as described below.

| Sample abbreviation | Species | Specimen form |
| --- | --- | --- |
| Human | *Homo sapiens* | Published PSMC |
| Gorilla | *Gorilla beringei* | Published PSMC |
| Sus-EUboar | *Sus scrofa* (wild boar from Europe) | Published reads |
| Sus-EUpig | *Sus scrofa* (domestic pig from Europe) | Published reads |
| Sus-CNboar | *Sus scrofa* (wild boar from China) | Published reads |
| Sus-CNpig | *Sus scrofa* (domestic pig from China) | Published reads |
| Aiptasia | *Aiptasia pallida* | Published reads |
| Acropora | *Acropora digitifera* | Published reads |
| Potato | *Solanum tuberosum* | Published reads |
| Agamb | *Anopheles gambiae* | Published reads |
| Pfalc-AS1 | *Plasmodium falciparum* (from Cambodia) | Published reads* |
| Pfalc-AS2 | *Plasmodium falciparum* (from Cambodia) | Published reads* |
| Tspir-US1 | *Trichinella spiralis* (from USA) | Biological sample |
| Tspir-US2 | *Trichinella spiralis* (from USA) | Biological sample |
| Tspir-CN1 | *Trichinella spiralis* (from China) | Biological sample |
| Tspir-CN2 | *Trichinella spiralis* (from China) | Biological sample |
| Smin | *Symbiodinium minutum* | Published reads |
| Pinf | *Phytophthora infestans* | Published reads |

**Table S2:**
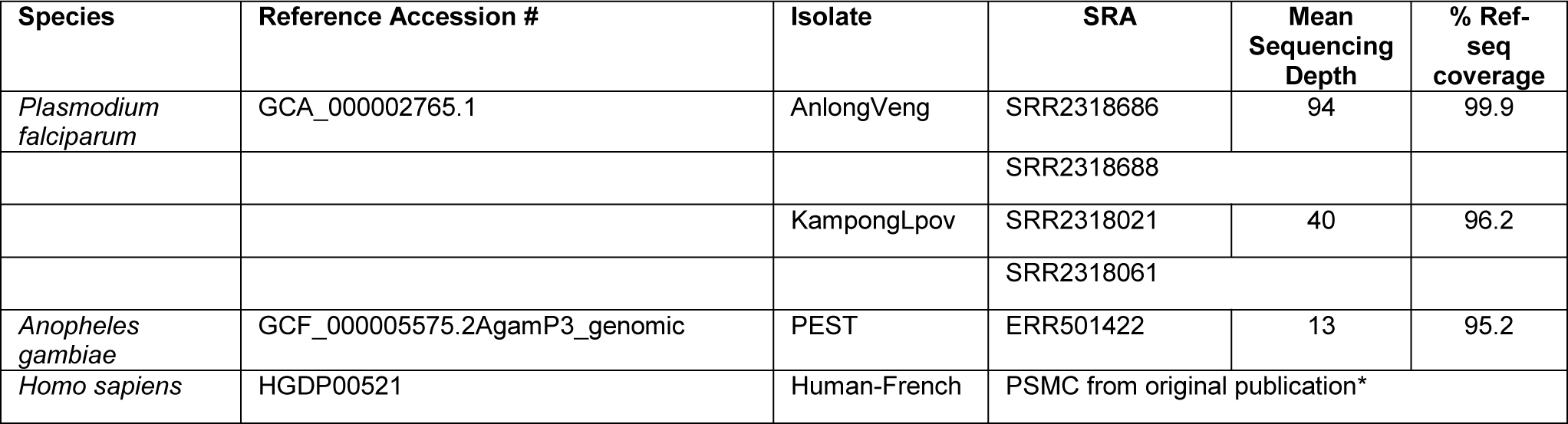
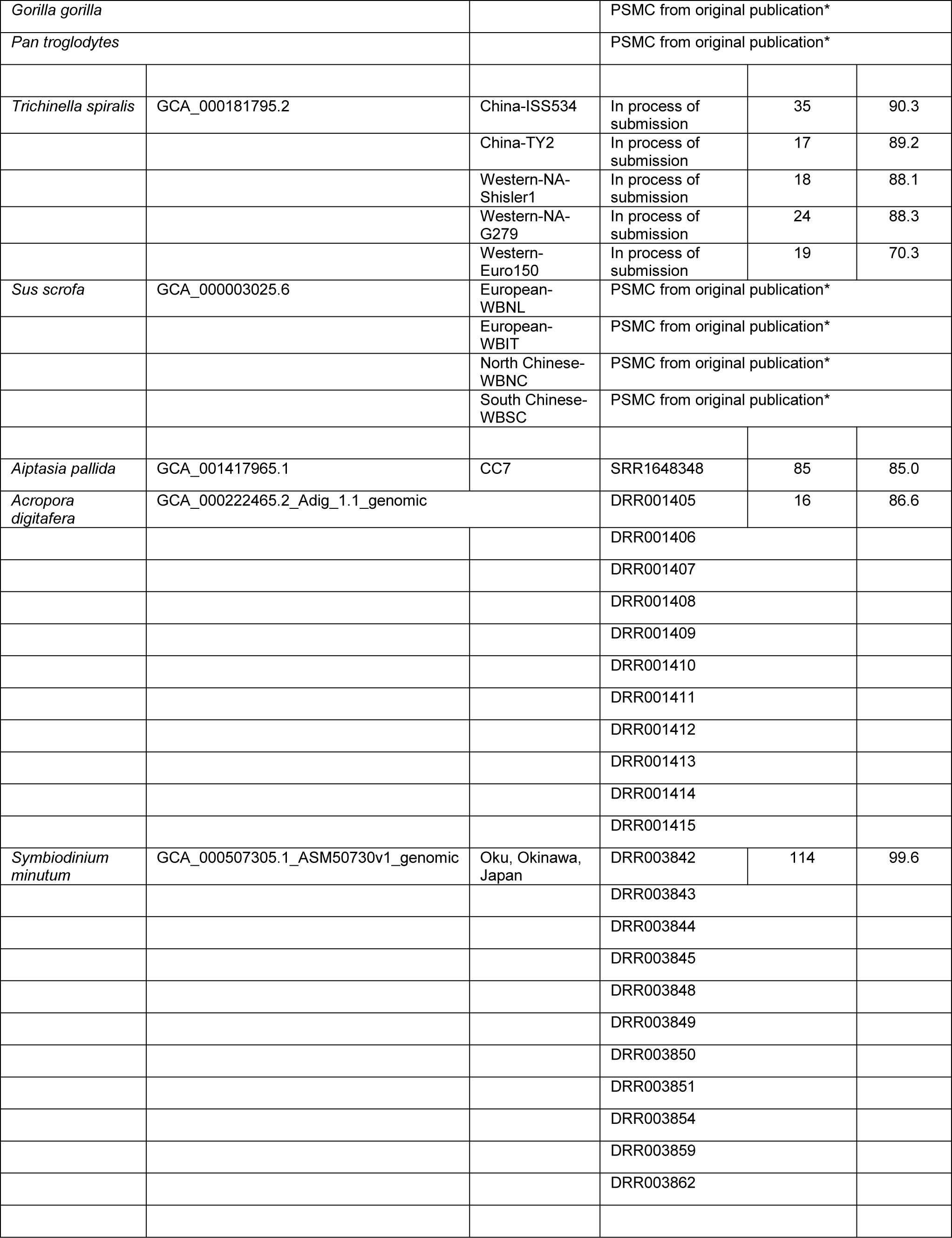
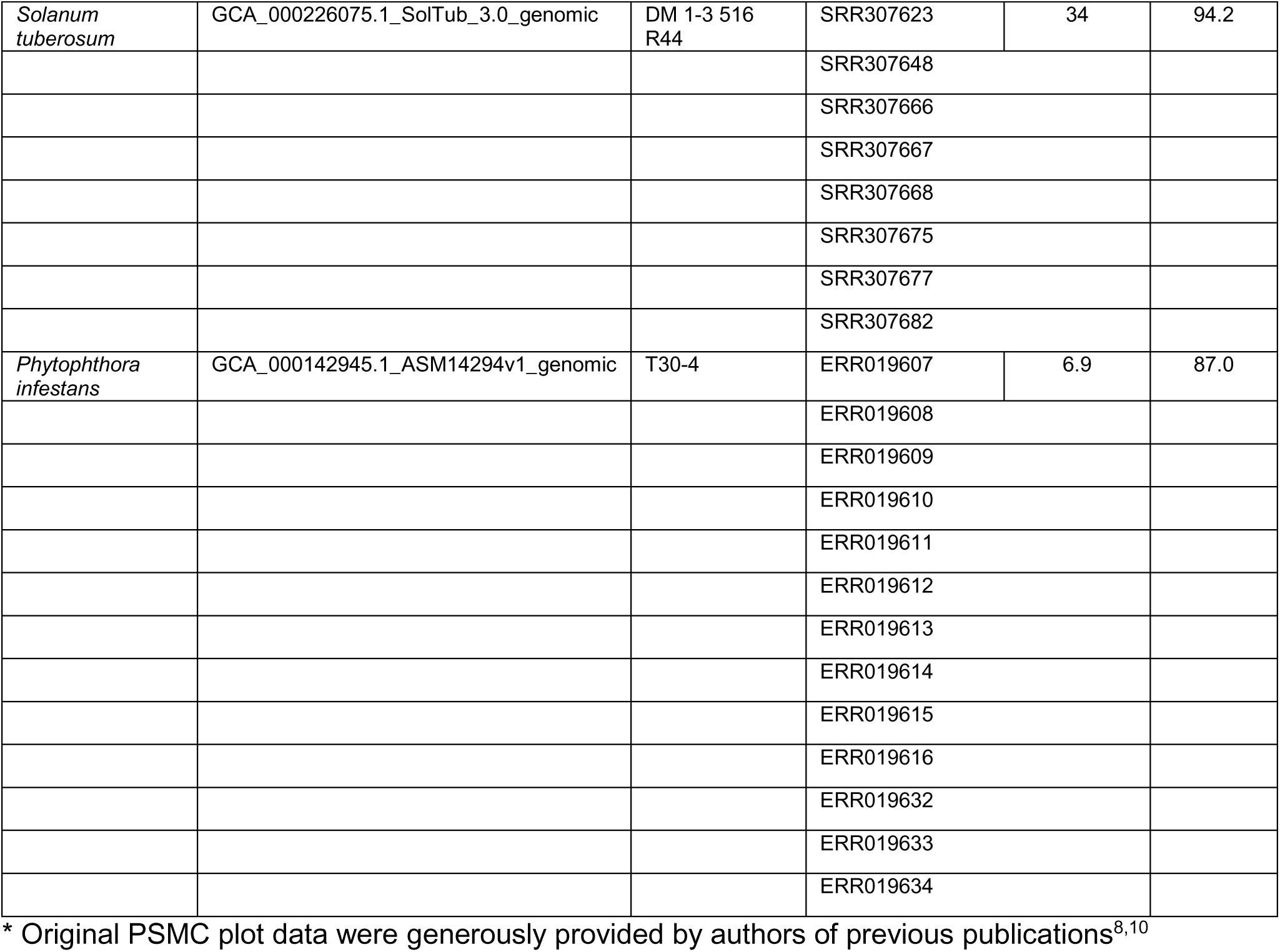
Summary of files used for genome assembly and heterozygote base- calling prior to PSMC analyses.

**Table S3:** Range of plausible generation times (in years) assumed for each symbiont genus.

| Species | Generation time (yr) |
| --- | --- |
| *A. gambiae* | 0.01 - 0.12 |
| *P. falciparum* | 0.3 - 0.8 |
| *T. spiralis* | 0.8 - 5.0 |
| *S. minutum* | 0.0 - 1.0 |
| *P. infestans* | 0.0 - 0.3 |

**Table S4:**
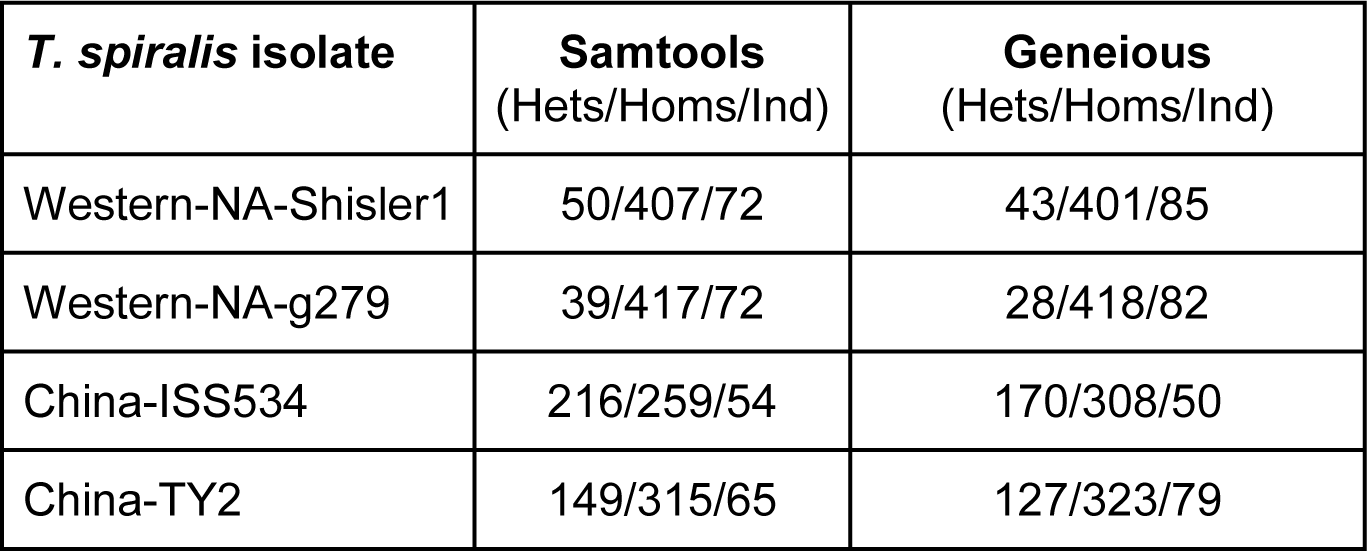
Estimated number of Heterozygotes (Hets)/Homozygotes (Homs)/Indeterminate (Ind) positions (x100,000) called by Samtools and Geneious in four isolates of *T. spiralis.*

| *T. spiralis* isolate | Samtools<br>(Hets/Homs/Ind) | Geneious<br>(Hets/Homs/Ind) |
| --- | --- | --- |
| Western-NA-Shisler1 | 50/407/72 | 43/401/85 |
| Western-NA-g279 | 39/417/72 | 28/418/82 |
| China-ISS534 | 216/259/54 | 170/308/50 |
| China-TY2 | 149/315/65 | 127/323/79 |

## References

1. Kuris, A. M. et al. Ecosystem energetic implications of parasite and free-living biomass in three estuaries. Nat. Lond. 454, 515–8 (2008).

2. Hechinger, R. F., Lafferty, K. D., Dobson, A. P., Brown, J. H. & Kuris, A. M. A common scaling rule for abundance, energetics, and production of parasitic and free-living species. Science 333, 445–448 (2011).

3. Hafner, M. S. & Nadler, S. A. Phylogenetic trees support the coevolution of parasites and their hosts. Nature 332, 258–259 (1988).

4. Page, R. D. M. & Charleston, M. A. Trees within trees: phylogeny and historical associations. Trends Ecol. Evol. 13, 356–359 (1998).

5. Hoberg, E. P. & Brooks, D. R. A macroevolutionary mosaic: episodic host-switching, geographical colonization and diversification in complex host–parasite systems. J. Biogeogr. 35, 1533–1550 (2008).

6. Li, H. & Durbin, R. Inference of human population history from individual whole-genome sequences. Nature 475, 493–496 (2011).

7. Liu, W. et al. Origin of the human malaria parasite *Plasmodium falciparum* in gorillas. Nature 467, 420–425 (2010).

8. Prado-Martinez, J. et al. Great ape genetic diversity and population history. Nature 499, 471–475 (2013).

9. Holt, R. A. et al. The genome sequence of the malaria mosquito *Anopheles gambiae*. Science 298, 129–149 (2002).

10. Joy, D. A. et al. Early origin and recent expansion of *Plasmodium falciparum*. Science 300, 318–321 (2003).

11. Consortium, T. A. gambiae 1000 G. Genetic diversity of the African malaria vector *Anopheles gambiae*. Nature 552, 96 (2017).

12. Rosenthal, B. M. et al. Human dispersal of *Trichinella spiralis* in domesticated pigs. Infect. Genet. Evol. 8, 799–805 (2008).

13. Zarlenga, D. S., Rosenthal, B. M., Rosa, G. L., Pozio, E. & Hoberg, E. P. Post-Miocene expansion, colonization, and host switching drove speciation among extant nematodes of the archaic genus *Trichinella*. Proc. Natl. Acad. Sci. 103, 7354–7359 (2006).

14. Groenen, M. A. M. et al. Analyses of pig genomes provide insight into porcine demography and evolution. Nature 491, 393–398 (2012).

15. Baumgarten, S. et al. The genome of *Aiptasia*, a sea anemone model for coral symbiosis. Proc. Natl. Acad. Sci. 112, 11893–11898 (2015).

16. Shoguchi, E. et al. Draft assembly of the *Symbiodinium minutum* nuclear genome reveals dinoflagellate gene structure. Curr. Biol. 23, 1399–1408 (2013).

17. Consortium, T. P. G. S. Genome sequence and analysis of the tuber crop potato. Nature 475, 189–195 (2011).

18. Haas, B. J. et al. Genome sequence and analysis of the Irish potato famine pathogen *Phytophthora infestans*. Nature 461, 393–398 (2009).

19. Lehmann, T., Hawley, W. A., Grebert, H. & Collins, F. H. The effective population size of *Anopheles gambiae* in Kenya: implications for population structure. Mol. Biol. Evol. 15, 264–276 (1998).

20. Hodges, T. K. et al. Large fluctuations in the effective population size of the malaria mosquito *Anopheles gambiae s.s.* during vector control cycle. Evol. Appl. 6, 1171–1183 (2013).

21. Rich, S. M., Licht, M. C., Hudson, R. R. & Ayala, F. J. Malaria’s Eve: Evidence of a recent population bottleneck throughout the world populations of *Plasmodium falciparum*. Proc. Natl. Acad. Sci. 95, 4425–4430 (1998).

22. Martin, F. N., Blair, J. E. & Coffey, M. D. A combined mitochondrial and nuclear multilocus phylogeny of the genus *Phytophthora*. Fungal Genet. Biol. 66, 19–32 (2014).

23. Zhou, X. et al. Whole-genome sequencing of the snub-nosed monkey provides insights into folivory and evolutionary history. Nat. Genet. 46, 1303–1310 (2014).

## References

1. Nadachowska-Brzyska, K., Burri, R., Smeds, L. & Ellegren, H. PSMC analysis of effective population sizes in molecular ecology and its application to black-and-white Ficedula flycatchers. Mol. Ecol. 25, 1058–1072 (2016).

2. Rich, S. M., Licht, M. C., Hudson, R. R. & Ayala, F. J. Malaria’s Eve: Evidence of a recent population bottleneck throughout the world populations of Plasmodium falciparum. Proc. Natl. Acad. Sci. 95, 4425–4430 (1998).

3. Volkman, S. K. et al. Recent origin of Plasmodium falciparum from a single progenitor. Science 293, 482–484 (2001).

4. Liu, W. et al. Origin of the human malaria parasite Plasmodium falciparum in gorillas. Nature 467, 420–425 (2010).

5. Escalante, A. A. & Ayala, F. J. Phylogeny of the malarial genus Plasmodium, derived from rRNA gene sequences. Proc. Natl. Acad. Sci. 91, 11373–11377 (1994).

6. Rich, S. M. et al. The origin of malignant malaria. Proc. Natl. Acad. Sci. 106, 14902–14907 (2009).

7. O’Loughlin, S. M. et al. Genomic signatures of population decline in the malaria mosquito Anopheles gambiae. Malar. J. 15, 182 (2016).

9. Manunza, A. et al. Romanian wild boars and Mangalitza pigs have a European ancestry and harbour genetic signatures compatible with past population bottlenecks. Sci. Rep. 6, 29913 (2016).

10. Groenen, M. A. M. et al. Analyses of pig genomes provide insight into porcine demography and evolution. Nature 491, 393–398 (2012).

11. Baker, A. C. Flexibility and specificity in coral-algal symbiosis: diversity, ecology, and biogeography of Symbiodinium. Annu. Rev. Ecol. Evol. Syst. 34, 661–689 (2003).

12. Rohling, E. J. Glacial conditions in the Red Sea. Paleoceanography 9, 653–660 (1994).

13. Biton, E., Gildor, H. & Peltier, W. R. Red Sea during the last glacial maximum: Implications for sea level reconstruction. Paleoceanography 23, PA1214 (2008).

14. Oppen, M. J. H. V., Willis, B. L., Vugt, H. W. J. A. V. & Miller, D. J. Examination of species boundaries in the Acropora cervicornis group (Scleractinia, Cnidaria) using nuclear DNA sequence analyses. Mol. Ecol. 9, 1363–1373 (2000).

15. Krasovec, M., Eyre-Walker, A., Sanchez-Ferandin, S. & Piganeau, G. Spontaneous mutation rate in the smallest photosynthetic eukaryotes. Mol. Biol. Evol. 34, 1770–1779 (2017).

16. Goodwin, S. B. The population genetics of. Phytopathology 87, 462–473 (1997).

17. Jaillon, O. et al. The grapevine genome sequence suggests ancestral hexaploidization in major angiosperm phyla. Nature 449, 463–467 (2007).

18. Li, H. & Durbin, R. Inference of human population history from individual whole-genome sequences. Nature 475, 493–496 (2011).

